# Temporal disaggregation through interval-integrated B-splines for the integrated analysis of trapping counts in ecology

**DOI:** 10.1101/2025.08.07.669113

**Authors:** Maxime Fajgenblat, Thomas Neyens

## Abstract

1. Passive trapping techniques such as pitfall and malaise traps probably constitute the most widely used methods for standardised surveys of invertebrate populations worldwide. These methods typically yield aggregated count data over multi-day trapping periods, often spanning several weeks, during which species activity (i.e. phenology) can vary. The analysis of trapping data collected over temporally misaligned sampling intervals is challenging, hampering the integrated analysis of historically available trapping datasets.
2. We introduce a temporal disaggregation approach using interval-integrated B-splines to analyse data collected over misaligned sampling intervals while accounting for phenological influences. We present computationally efficient Taylor series approximations for integrating exponentiated B-splines over sampling intervals. We further tailor our approach to typical trapping datasets by providing several extensions, including joint species distribution modelling.
3. Through simulations and cross-validation, we demonstrate that our approach of temporal disaggregation outperforms naive approaches and provides improved inference on phenology and other parameters of interest, such as inter-annual trends. The first-order Taylor approximation, which can be fit using regular software routines, properly accounts for heterogeneity in sampling duration and timing, while the second-order Taylor approximation and the exact model additionally allow for improved estimation of phenological patterns.
4. By applying this model to a large pitfall trapping dataset, spanning almost 50 years and over 10,000 trapping events in the Belgian province of Limburg, we illustrate how this approach can be used to reveal phenological, spatiotemporal and co-distributional patterns for 331 spider species.
5. The interval-integrated B-splines approach we present provides a convenient way to infer phenology and other ecological parameters from temporally aggregated count data obtained over misaligned sampling intervals, facilitating the integrated analysis of heterogeneously collected datasets to infer biodiversity trends.

## 1 Introduction

Passive trapping techniques rank among the most frequently used tools for studying invertebrate communities, offering standardised and low-effort sampling over extended periods. They typically involve placing a trapping device for a multi-day period, after which the captured individuals are retrieved for identification and counting. Pitfall trapping can be seen as the archetypical trapping technique and involves setting up a buried container with a preservation liquid flush with the soil, in which soil-dwelling animals get trapped and are being conserved (Greenslade, 1964). A wide variety of other trapping techniques are frequently used, including Malaise traps, flight interception traps and pan traps, each targeting different species guilds (Montgomery et al., 2021). Trapping methods have substantially advanced our understanding of invertebrate diversity and ecology, ranging from taxonomic discovery and uncovering ecological relationships (Bonte et al., 2003), to understanding the effects of management (Lawton et al., 1998; Rainio and Niemela, 2003; Longcore, 2003) and mapping insect biomass decline (Hallmann et al., 2017; Seibold et al., 2019). Since most trapping techniques are non-selective and therefore yield multi-species data, they have also received considerable attention in community ecology (e.g. Desender and Maelfait 1986). In this context, it is particularly illustrative that a pitfall trapping dataset has become the primary reference dataset used to illustrate and benchmark many of the developments in statistical community ecology over the past decades (e.g. ter Braak 1986; Peres-Neto et al. 2001; Hui et al. 2015; Popovic et al. 2019; van der Veen et al. 2021).

Although the cost-effectiveness of trapping techniques has contributed to their widespread use in ecological research, they have also attracted criticism. Numerous studies have documented discrepancies between trapping counts and actual species abundances (Luff, 1975), prompting the development of best practices that address both technical considerations and sampling design (e.g. Adis 1979; Ward et al. 2001; Brown and Matthews 2016; Engel et al. 2017). One key recommendation is that trapping counts should be viewed as proxies for species activity rather than direct measures of abundance (‘activity-abundance’; Heydemann 1953). Only catch totals obtained from trapping sessions that span at least a full reproductive cycle have been found to strongly correlate with actual abundances (Baars, 1979; Den Boer, 1979). These observations have led to the widely adopted recommendation that only trapping series covering a full year should be considered for quantitative analyses (Adis, 1979).

In most retrospectively available trapping datasets, trapping effort does not cover a full year and trapping events are asynchronous and misaligned. Discarding such datasets would be wasteful, especially given that the scarcity of biodiversity data already limits our capacity to assess conservation status, inform management decisions, and counteract shifting baselines (Kindsvater et al., 2018). Instead, Kindsvater et al. (2018) advocate the use of hierarchical statistical models for existing data sources to overcome the current biodiversity data crisis. Heterogeneity in sampling timing, along with phenological patterns (i.e. seasonal variation in species activity), and variation in sampling duration are two key challenges when analysing trapping data, both of which can strongly bias inference if left unaccounted for (Kotze et al., 2012). From a missing data perspective, properly accounting for trapping timing and duration is also required to mitigate the lack of a full-year sampling coverage and the non-random selection of sampling moments, assuming a missing at random mechanism (Rubin, 1976; Little and Rubin, 2019; Bowler et al., 2025). Yet, the vast majority of studies involving trapping data ignore heterogeneity in trapping timing and duration (Kotze et al., 2012).

Several statistical approaches have been put forward to model phenology in ecological data, ranging from mixture models and spectral decomposition methods to smoothing splines and Gaussian processes (Matechou et al., 2014; Bush et al., 2017; Strebel et al., 2014; Hefley et al., 2017). Their flexibility is particularly beneficial when analysing ecological data, in which complex functional relationships with time (e.g. caused by multivoltinism) are common. In combination with other modelling techniques, they offer opportunities to simultaneously mitigate biases when analysing opportunistically sampled or collated ecological data and to target specific research questions related to phenology (Lai, 2025). For instance, studying phenological change and its cascading effects is becoming particularly important as climate change is globally altering seasonal regimes (Parmesan, 2007; Hernández-Carrasco et al., 2025). The study of phenological change has long been dominated by descriptive metrics such as the first observation date, which have been shown to be very sensitive to population sizes, sampling effort and extreme events (Miller-Rushing et al., 2008; Van Strien et al., 2008; Moussus et al., 2010). Approaches such as smoothing splines offer opportunities to overcome these limitations, and have therefore been recommended instead (Moussus et al., 2010).

A major complication in modelling phenological patterns using trapping data arises from the aggregated nature of trapping counts: they are only available at the resolution of the sampling interval. With the exception of two applications related to spatial downscaling in species distribution modelling (Keil et al., 2013; Murphy et al., 2023), the analysis of aggregated data has received remarkably little attention in the field of ecology, in contrast to several other scientific disciplines. In hydrology and climatology, for instance, these methods are routinely used to downscale coarse climate model outputs, transforming, for example, daily precipitation data into hourly series to capture diurnal cycles and extreme events (Connolly et al., 1998; Zabel and Poschlod, 2023). In econometrics, they underpin mixed-data sampling (MIDAS) regressions that link low-frequency macroeconomic indicators with high-frequency financial data to enhance forecasting accuracy (Ghysels et al., 2004). In epidemiology, spatial disaggregation has emerged as a promising approach for spatial downscaling as well as for the combined analysis of multi-resolution datasets, and gained much attention from a methodological and applied point of view (Wakefield and Salway, 2001; Diggle et al., 2013; Weiss et al., 2019; Johnson et al., 2019; Rutten et al., 2025).

In this paper, we explore the data-generating process that leads to aggregate counts in trapping data, while accounting for phenological patterns, to derive a B-spline based modelling approach that addresses heterogeneities in sampling duration and timing. We present computationally efficient approximations and we evaluate model performance through a simulation study. We demonstrate how phenological change can be inferred and how a spatio-temporal joint species distribution model can be constructed for temporally aggregated data. Throughout our applications, we use an extensive pitfall trapping dataset collected over asynchronous time intervals, spanning a total period of almost 50 years.

## 2 Material and methods

### 2.1 Temporal disaggregation using interval-integrated B-splines

#### 2.1.1 Derivation

Let *Y*_*i*_ denote the random variable representing the number of individuals observed during sampling event *i* = 1, 2, …, *n*. Since sampling events typically span multiple days, the random variable *Y*_*i*_ can be decomposed into the sum of (unobserved) daily counts: 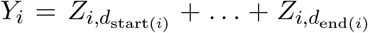, where *Z*_*i,d*_ denotes the random variable for the number of individuals during day *d* for the *i*-th sampling event. We assume that *Z*_*i,d*_ follows a Poisson distribution:

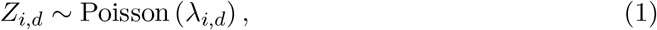

where *λ*_*i,d*_ is the expected number of sampled individuals during day *d* for the *i*-th sampling event. Following the generalised additive model (GAM) framework (Hastie and Tibshirani, 1986; Wood, 2017), we model the linear predictor as the sum of a parametric samplespecific linear predictor part *µ*_*i*_ and a non-parametric day-specific phenological part *f* (*d*):

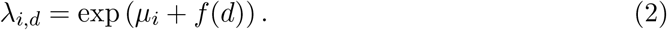

We assume the phenological activity function *f* (*d*) can be described using cubic B-splines, as the weighted sum of an appropriate number of *k* basis functions:

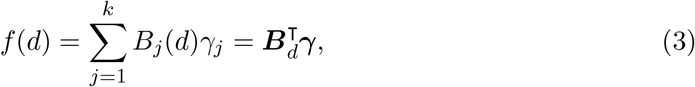

where *B*_*j*_(*d*) is the *j*-th basis function and *γ*_*j*_ is the corresponding basis function weight, and where 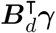 constitutes a dense inner product notation. When considering phenomena that span a full year, the use of cyclic B-splines is advisable to ensure seasonal patterns match at the turn of the year.

Since the daily outcomes are unobserved, we now return to the linear predictor for aggregate outcomes. Using an identity link function (e.g. for Gaussian responses), the phenological part of the aggregated linear predictor would simply reduce to the Riemann integral of the phenological curve over the sampling interval (Supplementary Information, Section S1.1). Conveniently, this can be achieved by supplying a design matrix of *interval-integrated basis functions* to a GAM model instead of a conventional basis function design matrix. Such a design matrix can easily be pre-computed prior to estimation by evaluating and summing the daily basis functions over the days of the sampling interval of sampling event *i* (Supplementary Information, Section S1.1).

For Poisson responses, the aggregate count *Y*_*i*_ can be expressed in terms of the daily expected counts 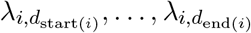, assuming conditional independence among counts of the individual days comprised in sampling event *i*:

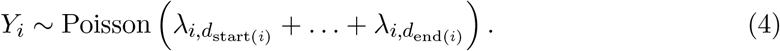

The corresponding linear predictor therefore reduces to:

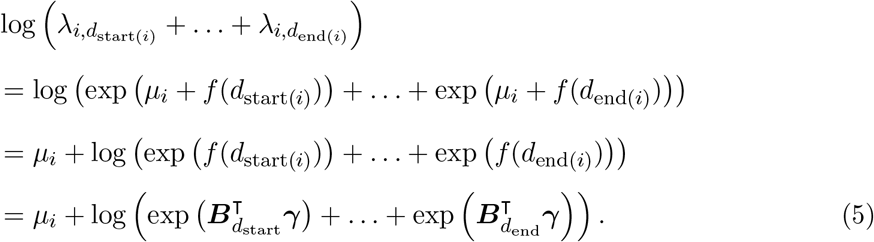

In contrast to the Gaussian case, this expression cannot be further simplified by substitution of pre-computed interval-integrated basis functions, and the computation of the sum of daily exponentials for each sampling event can be prohibitive for larger datasets (cf. simulations below). A computationally efficient approximation of the interval-integrated exponentiated B-splines can, however, be achieved through a Taylor series expansion around the midpoint 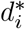 of the sampling interval. The first-order Taylor approximation simplifies to a conventional offset model (see Supplementary Information, Section S1.2 for a full derivation):

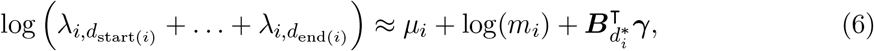

where *m*_*i*_ is the duration of the sampling interval. This approach is similar to the one proposed by Kotze et al. (2012), with the exception that phenology is now modelled continuously through smoothing splines instead of discrete factor levels that correspond to non-overlapping sampling intervals. The present approach, therefore, enables the estimation of more flexible and realistic phenological patterns, and allows for arbitrarily overlapping sampling intervals, further relaxing dataset requirements and facilitating broad-scale data-integration efforts. While this first-order approximation can readily be implemented using conventional GAM fitting routines such as the gam function of the mgcv package in R (Wood, 2017), it implies a constant phenology over the course of the sampling interval. The second-order Taylor approximation includes an additional term that accounts for phenological variation during the sampling interval through the first and second derivative of the phenological function (see Supplementary Information, Section S1.2 for a full derivation):

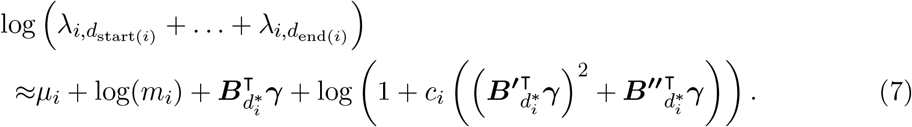

This linear predictor is computationally cheap and does not require the estimation of additional parameters, because 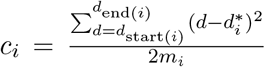 is a variance-like constant which can be computed ahead of model estimation, and because of the ease of differentiating\ B-splines using the derivative basis function evaluations 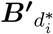 and 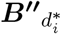, which can also be pre-computed for each sampling event *i*.

#### 2.1.2 Regularised phenological activity functions

Choosing the number of basis functions is known to be critical when using B-splines, rendering them sensitive to over- and underfitting (Eilers and Marx, 1996). The use of wigglyness penalties in penalised B-splines (P-splines) allows the user to overspecify the number of basis functions, while constraining the basis function weights from being too wiggly (Eilers and Marx, 1996). In the frequentist setting, optimal penalties are typically determined through prediction error minimizing approaches (such as generalised cross-validation or AIC), or through likelihood-based approaches. Bayesian P-splines replace conventional penalties by random walk priors to regularise basis function weights and to enforce smoothness, enabling simultaneous estimation and uncertainty propagation of both the smooth function and smoothing parameters (Lang and Brezger, 2004; Miller, 2025). In our models, we specifically use B-splines projected Gaussian process priors, which have been shown to outperform (Bayesian) P-splines and match the performance of Gaussian processes, while being computationally cheaper (Monod et al., 2023). Instead of random walk priors, B-splines projected Gaussian processes feature a more powerful and flexible Gaussian process prior on the basis function weights (Monod et al., 2023). Since we consider cyclic B-splines to ensure seasonal patterns match at the turn of the year, we specifically use a periodic kernel *C* as covariance function (Rasmussen and Williams, 2006):

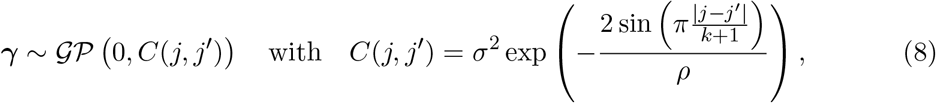

where *j* is the basis function index (cf. Eq. 3), *k* the total number of basis functions (cf. Eq. 3; with *k* + 1 being the kernel’s periodicity), *σ* the phenological activity function’s scale (‘amplitude’) parameter, and *ρ* the phenological activity function’s length scale (‘turnover’) parameter. Conveniently, the use of Bayesian priors to regularise the basis function weights does not interfere with the structure of the temporal disaggregation models presented in Eqs. 5-7. It only requires setting up a sufficiently large number of equally spaced basis functions for the regularization to act on.

#### 2.1.3 Yearly varying phenological activity functions

As phenological patterns can vary over years, it can be important to allow for yearly varying phenological activity functions when analysing multi-year data. Data sparseness, however, will often preclude the estimation of yearly independent phenological activity functions. We therefore propose the following efficient yet flexible approach to estimate the yearly basis function weights ***γ***_***t***_, by allowing for linearly weighted deviations from the historical basis function weights the over time:

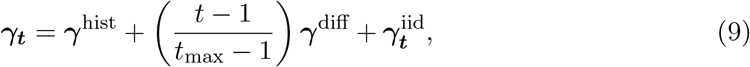

where ***γ***^hist^ is a set of historical basis function weights (corresponding to *t* = 1), ***γ***^diff^ is a set of contemporary basis function weight differences (capturing the phenological change between *t* = *t*_max_ and *t* = 1), and 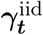 is a set of yearly (i.i.d.) normal deviations from the linearly changing basis function weights over time. This approach allows the estimated phenology to change linearly and directionally over years, while allowing for yearly non-directional deviations. This specific model structure can be simplified or extended depending on a priori knowledge on the functional relationship of phenological change, or based on research questions at hand. Similarly to the regularization of the basis function weights (cf. Section 2.1.2), the use of yearly varying phenological activity functions does not interfere with the structure of the temporal disaggregation models presented in Eqs. 5-7. It can, however, slow down computation by requiring indexing to apply the appropriate set of basis function weights based on each sampling event’s year, precluding efficient vectorization across all sampling events.

### 2.2 Towards a comprehensive modelling approach for trapping data

In the following, we provide several model extensions to further accommodate the characteristics of typical trapping datasets.

#### 2.2.1 Accounting for overdispersed counts

Ecological count data is typically overdispersed, leading to deviations from the Poisson distribution’s mean–variance relationship (Ver Hoef and Boveng, 2007). To accommodate this, we extend our model to a negative binomial likelihood (O’Hara and Kotze, 2010). Since the negative binomial distribution can be derived as a Poisson-Gamma mixture (Hilbe, 2011), it can be shown that the aggregate count 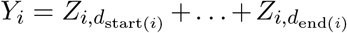 also follows a negative binomial distribution with overdispersion parameter *ϕ* (see derivation in the Supplementary Information, Section S1.3):

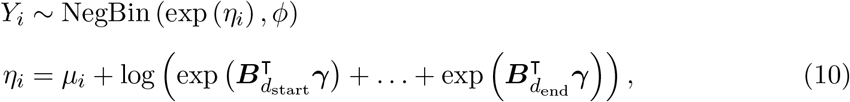

or alternative specifications in which the linear predictor is replaced by the Taylor approximations presented in Eq. 6 and Eq. 7.

#### 2.2.2. Extending the linear predictor for trapping data

In Eq. 2, we introduced *µ*_*i*_ as the part of the linear predictor that describes variation among samples unrelated to the phenology. Although the precise structure of the linear predictor should be tailored to the application at hand, we here propose a flexible and generic structure for ecological applications, that accounts for variation induced by a set of covariates, space and time:

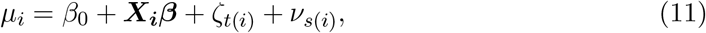

where *β*_0_ is the model intercept, ***X*** is a covariate design matrix (e.g. sampling method or attractant used, or distributional predictors for the sampled site *s*(*i*)), ***β*** is a vector of corresponding regression coefficients, *ζ*_*t*(*i*)_ is a temporal random effect for the sampled year *t*, and *ν*_*s*(*i*),*j*_ is a spatial random effect for the sampled site *s*. The temporal random effects *ζ*_*t*_ are modelled using exact Gaussian processes (Rasmussen and Williams, 2006), while the spatial random effects *ν*_*s*_ consist of a spatially structured (using B-splines projected Gaussian processes; Monod et al. 2023) and a spatially unstructured (normal i.i.d. random effects) part. We provide full methodological detail for these model components in the Supplementary Information (Section S1.4).

#### 2.2.3 Joint species distribution modelling

Since non-selective trapping methods typically yield multi-species data, we embed the outlined temporal disaggregation structure in a joint species distribution model. By parsimoniously sharing information across species, this embedding can be expected to improve species-specific inference, especially benefiting rare species (Ovaskainen and Soininen, 2011; Warton et al., 2015). Since the exact linear predictor structure (Eq. 5) is too computationally demanding when embedded in a joint species distribution model, we only consider the computationally efficient approximations.

The aggregate counts *Y*_*i,l*_ for sampling event *i* and species *l* are modelled simultaneously in a single model as follows:

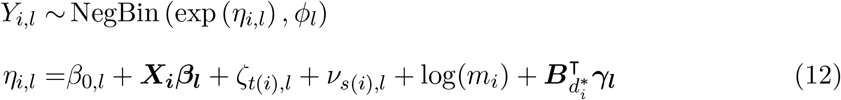

or with a second-order Taylor approximation as presented in Eq. 7. The interspecific exchange of information is achieved at three levels in this model.

First, the intercepts *β*_0,*l*_ and regression coefficients ***β***_***l***_ are each estimated hierarchically, as normally distributed random effects with a certain spread around the community-level mean coefficient value (Zipkin et al., 2009). This approach has been shown to improve precision for rare species (Zipkin et al., 2009), and provides the basis for further extensions in pooling information across species (not considered here), e.g. by accounting for phylogenetic relatedness and trait influences (Ovaskainen et al., 2017).

Second, the spatial random effects *ν*_*s*(*i*),*l*_ are estimated using a latent factor approach, to condensate distributional patterns in a limited number of latent dimensions *q*_max_:

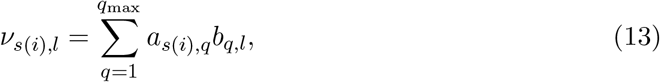

where *a*_*s*(*i*),*q*_ and *b*_*q,l*_ represent site and species loadings, respectively. The set of site loadings *a*_·,*q*_ per latent dimension *q* are modelled in the same fashion as the single-species spatial random effects in Eq. 11, using Gaussian processes. Identifiability is achieved through a multiplicative gamma process shrinkage prior on the species loadings, as described in Bhattacharya and Dunson (2011) and as implemented in state of the art joint species distribution models (Ovaskainen et al., 2017; Tikhonov et al., 2020; Norberg et al., 2019).

Third, the species-specific phenological basis function weights ***γ***_***l***_ are estimated following a hierarchical generalised additive model (HGAM; Pedersen et al. 2019) framework:

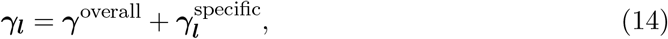

where ***γ***^overall^ is the common phenological pattern shared by all species and 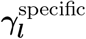 captures species-specific deviations from the common phenological pattern. Assuming many members of the species community share a largely similar phenological activity pattern, this structure improves phenology estimation, especially for rare species, while allowing other species to deviate from the overall pattern based on available data. Both ***γ***^overall^ and 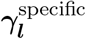 are estimated using the B-splines projected Gaussian process priors, with distinct scale and length scale parameters across both levels and species (type GI, *sensu* Pedersen *et al.* 2019).

In addition to the dispersion parameter ϕ_*l*_, we estimate the temporal random effects *ζ*_*t*(*i*),*l*_ in a strictly species-specific fashion (i.e. without any pooling of information across species) to avoid any generalization bias.

### 2.3 Implementation

We implemented the modelling approach using the probabilistic programming language Stan (Carpenter et al., 2017). Stan performs Bayesian inference by means of dynamic Hamiltonian Monte Carlo (HMC), a gradient-based Markov chain Monte Carlo (MCMC) sampler (Betancourt, 2018). We use the Pathfinder algorithm, a recently developed quasi-Newton variational inference method that has been shown to quickly reach the high probability region of complex target distributions, to initialise MCMC chains (Zhang et al., 2022). To fit the model, we use the ‘CmdStanR’ package as an interface to Stan v2.36.0, in R v4.3.1 (R Core Team, 2023). We chose weakly informative priors for all parameters (see Supplementary Information, Section S1.5).

The R- and Stan-code related to all outlined analyses is available on GitHub through https://anonymous.4open.science/r/trappingmodel-750A/. We present four different Stan model scripts, differing in the extensions outlined in Sections 2.1–2.2 that are implemented (Table 1). Each of these scripts corresponds to up to eight possible submodels through the ability to consider among four disaggregation approaches and two model likelihoods, which can be specified as part of the provided data. In addition to the first- and second-order Taylor approximations and the exact disaggregation approaches discussed in Section 2.1, most models also allow for a naïve approach that only features a logarithmic offset to account for variation in sampling duration, but that does not feature a phenological model component. The model that does not feature an extended linear predictor (`TDM_simple.stan`) only offers the possibility to account for covariates by supplying a design matrix, in addition to the intercept and the temporal disaggregation term.

**Table 1:** Overview of the four temporal disaggregation models and their features used throughout this study, implemented as Stan scripts.

| Model script name (*.stan) | TDM_simple | TDM_flexible | TDM_yearlyphen | TDM_multispecies |
| --- | --- | --- | --- | --- |
| Disaggregation approach (Section 2.1) | Naïve<br>Taylor 1<br>Taylor 2<br>Exact | Naïve<br>Taylor 1<br>Taylor 2<br>Exact | Naïve<br>Taylor 1<br>Taylor 2<br>Exact | Taylor 1 |
| Regularised phenological activity function (Section 2.1.2) | ✓ | ✓ | ✓ | ✓ |
| Yearly phenological activity function (Section 2.1.3) | ✗ | ✗ | ✓ | ✗ |
| Implemented model likelihoods (Section 2.2.1) | Poisson<br>Neg. binomial | Poisson<br>Neg. binomial | Poisson<br>Neg. binomial | Neg. binomial |
| Extended linear predictor (Section 2.2.2) | ✗ | ✓ | ✓ | ✓ |
| Multi-species outcomes (Section 2.2.3) | ✗ | ✗ | ✗ | ✓ |

We run eight MCMC chains using 1,000 iterations each, of which the first 500 were discarded as warm-up, unless otherwise specified. We assessed model convergence both visually by means of trace plots (for a random subset of parameters) and numerically by means of effective sample sizes, divergent transitions and the Potential Scale Reduction Factor (Vehtari et al., 2021), for which all runs had *R <* 1.1.

### 2.4 Simulation study

We performed a simulation study to assess the performance of the outlined temporal disaggregation models to infer phenology and secondary model parameters. We assess whether model performance varies with respect to (1) the duration of sampling intervals; and (2) the rate at which phenological activity varies throughout the year. In each run, we simulate 1000 trapping events spanning either 15 or 30 days (reflecting different sampling interval durations), randomly spread between day 1 and day 365 of the year. In each run, we simulate a randomly drawn phenological activity function using cyclic B-splines with scale 5, and with either 8 or 16 basis functions (reflecting different rates at which phenological activity vary). We use a model intercept of −4 and we simulate a covariate gradient using a uniform distribution ranging from 0 to 1 across the 1000 sampling events, along which the linear predictor for abundance varies with a slope of −1. This slope can be thought of a negative abundance trend one might want to infer, as a secondary inferential target in addition to phenological activity patterns. Together, the aforementioned intercept, covariate gradient and corresponding slope, as well as the phenological activity curve represent the expected number of trapped individuals during a single day (on the logarithmic scale), used to draw the realised number of trapped individuals using a Poisson distribution. The daily trapping counts are subsequently summed to achieve the observed, aggregate trapping count. We explore the two simulation settings in a full factorial way, yielding four scenarios, each replicated 50 times.

We assume phenology to be the only driver of temporal variation during a single sampling event and that the number of trapped individuals is ignorable with respect to local populations sizes (a common assumption in trapping studies). We do not account for a varying trapping intensity throughout trapping intervals. We restrict our simulations to the assessment of the main novelty of the present study, i.e. the temporal disaggregation model. We therefore focus on the single-species implementation only, since the considered joint species distribution modelling approaches have been validated and evaluated elsewhere (e.g. Norberg et al. 2019).

We evaluate the eight possible submodels for the model script `TDM_simple.stan` (Table 1). We consider the following performance metrics: (1) coverage of the phenological activity function (fraction of days for which the true phenological activity value falls within the 95% credible interval); (2) mean absolute error of the phenological activity function (mean absolute deviation of the posterior mean phenological activity function compared to the true function, averaged over days); (3) trend uncertainty width (length of the 95% credible interval of the posterior trend slope); (4) mean absolute error of the trend slope (mean absolute deviation of the posterior mean trend slope compared to the true value); and (5) model run time (duration of the slowest MCMC chain).

### 2.5 Application: spider pitfall trapping data in Limburg

We apply our temporal disaggregation approach to a large pitfall trapping dataset collected over a 49-year period (1976-2024) in the province of Limburg, in the north-east of Belgium. The dataset is managed by the Limburgse Koepel voor Natuurstudie (LIKONA) and has largely been collected by expert volunteers and professionals of this organization.

Trapping events were only retained if the exact deployment date, retrieval date and spatial coordinates were known. Additionally, trapping events with a duration exceeding 60 days were discarded. Species that were detected in less than 10 trapping events, were also discarded from the dataset. Our analysis features 12,451 trapping events, covering 1,346 distinct locations across Limburg (Fig. S1) and 331 spider species. In total, 704 ‘trap-years’ were performed. Sampling intervals and durations are highly heterogeneous across sampling events and years (Fig. S2-3).

Although the majority of trapping events were performed using conventional pitfall traps, three variants were sometimes used: ramp traps, pyramid pitfall traps, subterranean pitfall traps, aquatic traps and cavity traps. These different variants were included in the analysis as covariates using dummy coding, with conventional pitfall traps as reference level, allowing for differential affinities among species (cf. Section 2.2.4). Further details regarding the trap construction and preservation fluid were not documented in the dataset, but interviews with data collectors revealed that the majority of traps consisted of glass cups, in addition to some occasions when plastic cups were used. Both glass and plastic cups had a 9 cm diameter. The type of preservation liquid administered to the cups changed over time: picric acid was used until the mid 1980’s, after which a 2-4% formaldehyde solution was used. This was gradually replaced by alternatives such as polypropylene glycol. Since we do not have reliable data on these trap characteristics, we did not correct for them in the analysis.

We use 15 equally spaced cyclic basis functions for modelling the phenological activity functions (through B-splines projected Gaussian processes), assuming this is largely sufficient to capture fine-grained phenological patterns. To account for spatial patterns across the sampling locations, and to be able to map spatial patterns for the entire province of Limburg, we link all pitfall trapping locations to a 1 × 1 km grid defined over the study areas. We use 15 × 15 equally spaced basis functions for modelling spatial patterns across the grid cells (through B-splines projected 2D Gaussian processes). In addition to the the pitfall trapping variant information, we included land cover fractions of 14 land cover types for each of the grid cell as predictor, based on the Biological Valuation Map (De Saeger et al., 2020). The dominant land cover type, arable land, was omitted to avoid overspecification due to compositionality.

We only consider the negative binomial model likelihood for this real-data application because preliminary explorations revealed substantial overdispersion.

#### 2.5.1 Inferring multi-decadal phenological change in soil-dwelling spiders

To test the ability of the outlined approaches to account for and to infer phenology and phenological change over years using temporally aggregated data, we applied the model script `TDM_yearlyphen.stan` (Table 1) in a species-by-species fashion to the spider dataset. For each species, each of the four temporal disaggregation methods was fitted. We used the leave-one-out cross-validation information criterion (LOOIC; Vehtari et al. 2017) to compare predictive performance. For each year and at each posterior draw (i.e. fully propagating uncertainty), we derived the day of the year at which the estimated phenological activity function reaches the highest value, i.e. the peak activity day. We quantify the evidence of change in peak activity days by regressing these values against the year through ordinary least squares (OLS), yielding the slope of phenological change over the study period at each posterior draw. We compute the 1-Wasserstein distance, also known as the earth mover’s distance (Panaretos and Zemel, 2019), to compare the posterior slopes obtained from the first- and second-order Taylor approximations to the ones obtained from the exact model, which we consider to be the reference model. Higher 1-Wasserstein distance indicate inferior inferential quality, accounting for both bias and variance, by quantifying the minimal effort to reconfigure one posterior distribution into another (Panaretos and Zemel, 2019). We do not include the naïve model in these specific comparisons as this approach does not contain any model component to infer phenology. We restricted this analysis to the 50 most abundantly caught species.

#### 2.5.2 Joint species distribution modelling of soil-dwelling spider communities

To demonstrate how to perform joint species distribution modelling and model-based ordination using temporally aggregated data, we applied the model script `TDM_multispecies.stan` (Table 1) to the spider dataset. We consider all 331 species, and we specified 10 latent dimensions for the latent factor approach. We specifically chose the first-order Taylor approximation as the exact linear predictor structure was too computationally demanding for the multi-species setting, and as this approximation turned out to be sufficient when one wants to acknowledge the influence of phenology, but when the main interest lies in other inferential targets, as will be demonstrated below.

## 3 Results

### 3.1 Simulation study

Our simulation study shows that failing to address phenological patterns in aggregated ecological count data by merely accounting for sampling duration can negatively impact inference on quantities of interest, such as simulated inter-annual trends considered in our simulation study (Fig. 1). All evaluated models that account for for phenology strongly outperform the naïve model and perform similarly with respect to estimating such secondary quantities of interest. When failing to properly account for temporal aggregation in the face of phenological patterns, the negative binomial model likelihood is able to capture the overdispersion resulting from the process, while the use of a Poisson model likelihood leads to an underestimation of uncertainty. Our simulations show that the performance of the first-order Taylor approximation matches the second-order Taylor approximation and the exact model when inferring quantities of interest unrelated to phenology (Fig. 1).

**Figure 1.**
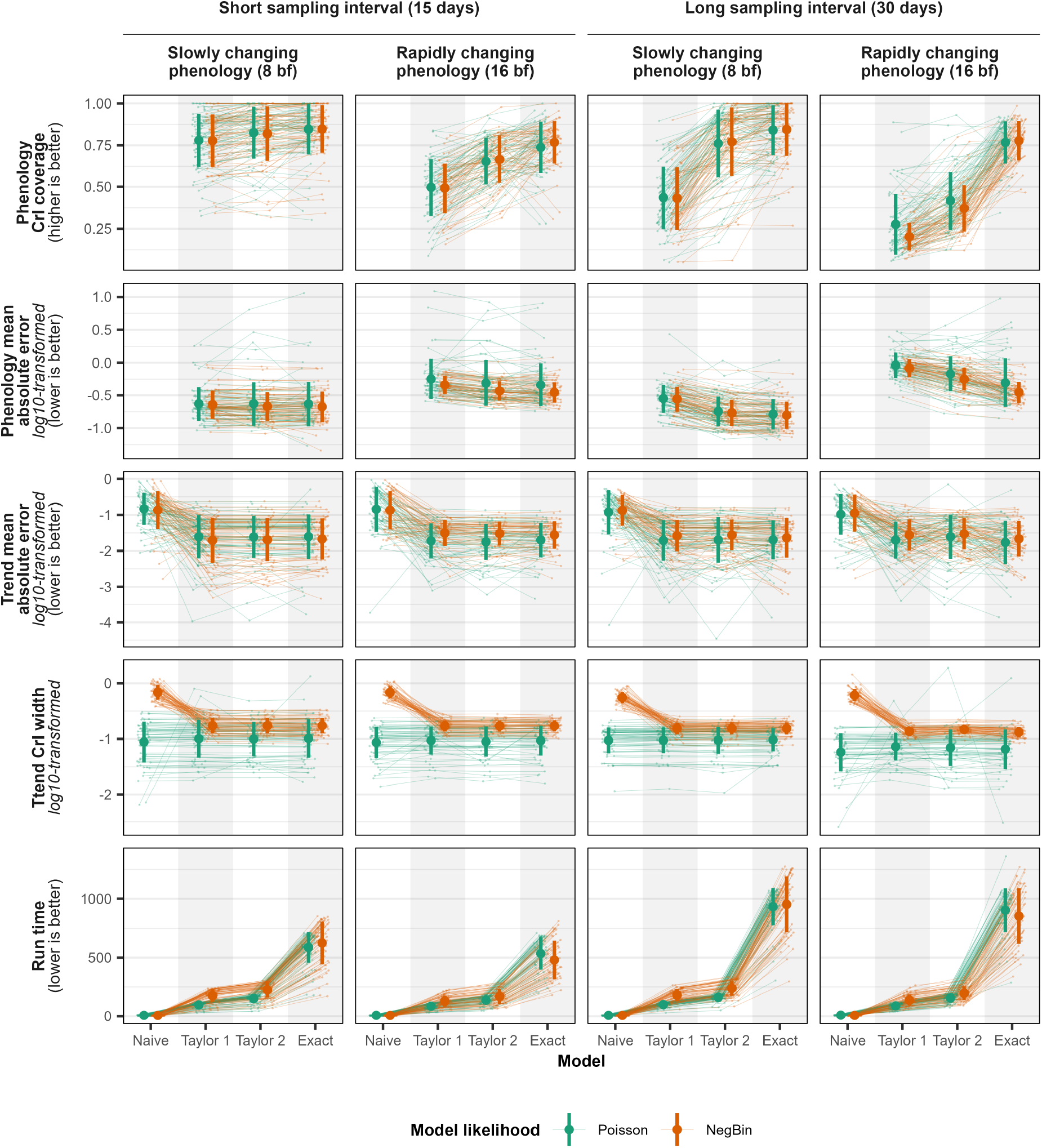
Performance of the eight considered models to infer phenology and a trend parameter.

The exact model outperforms the approximations for inferring phenological patterns, both in terms of credible interval coverages and mean absolute errors. In turn, the second-order Taylor approximation outperforms the first-order Taylor approximation in the vast majority of simulations. The gains of opting for a more complex model structure, however depend on the sampling interval duration as well as the rate of phenological change. These gains are highest in scenarios with a rapidly changing phenology with long sampling intervals. In scenarios with slowly changing phenology and long sampling intervals, the performance of the second-order Taylor approximation is almost equivalent to the one of the exact model. In scenarios with a slowly changing phenology and short sampling intervals, the performance of the three approaches is very similar (Fig. 1).

The computational cost varies widely across models. It is lowest for the naïve model, higher for the first- and second-order Taylor approximation (with both being very similar), and exceedingly higher for the exact model. For computational cost for the exact model increases along with the sampling duration.

### 3.2 Inferring multi-decadal phenological change in soil-dwelling spiders

The overall predictive performance, as quantified through LOOIC, is markedly higher for the first- and second-order Taylor approximations as well as for the exact model (Fig. 2). Our results suggest that the first-order Taylor approximation is sufficient to optimise predictive performance, as the second-order Taylor approximation and the exact model no not offer any detectable advantage in terms of LOOIC (Fig. 2).

**Figure 2.**
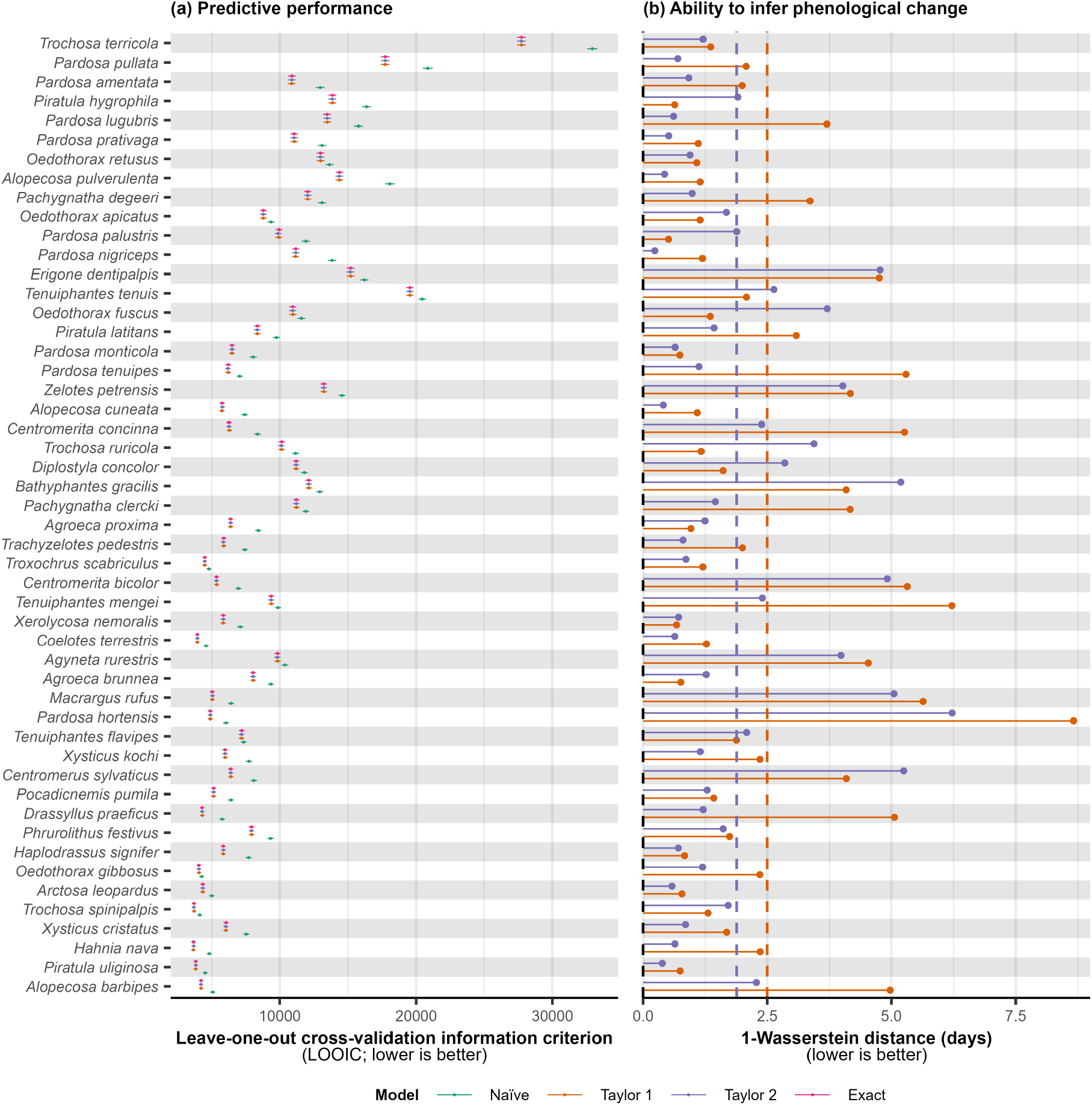
Model performance comparison, considering the 50 most abundantly caught spider species included in the case study. (a) Predictive performance, as measured by the leave-one-out information criterion (LOOIC) for the four possible models. LOOIC values are depicted by points, while error bars represent standard error values. LOOIC values should only be compared within species, and not across species. (b) Ability to infer phenological change, as measured by the 1-Wasserstein distance. The first- and second order Taylor approximations are benchmarked against the exact model. The naïve model does not allow inference on phenological change, and is not included in this comparison. The dashed lines represent averages across all species

When inferring phenological change, the second-order Taylor approximation outperforms the first-order Taylor approximation, assuming the phenological change inferred by the exact model to be the closest to the truth (Fig. 2). The average 1-Wasserstein distance for the first-order Taylor approximation equals 2.5 days, while it equals 1.9 days for the second-order Taylor approximation, with a significant difference between both methods (Wilcoxon signed-rank test; *V* = 956; *p* = 0.006).

An in-depth exploration of the actually inferred phenological patterns and multidecadal change thereof is beyond the scope of this publication. We therefore restrict ourselves to outlining the results for the four most abundantly caught species in the datasets: *Trochosa terricola, Pardosa pullata, Pardosa amentata* and *Piratula hygrophila*. These species displayed a posterior mean phenological advancement by 12.8 (95% CrI [0.5,32.7]), 33.4 (95% CrI [17.2,52.0]), 18.2 (95% CrI [-1.3,38.8]) and 28.1 (95% CrI [1.8,32.7]) days, respectively, over the 49-year study period, as illustrated in Fig. 3.

**Figure 3.**
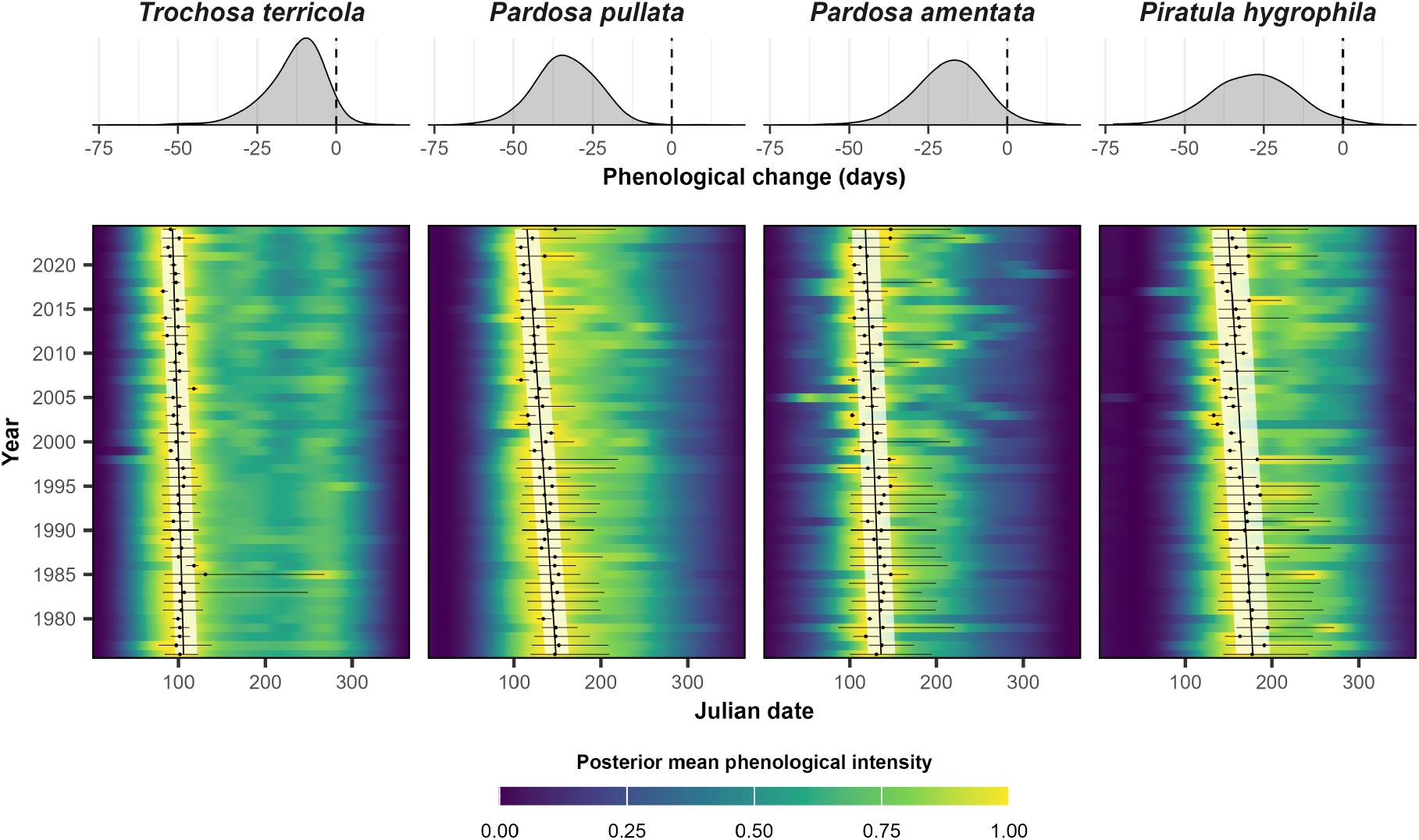
Estimated yearly phenological activity patterns and change thereof for the four most abundantly caught spider species included in the case study, obtained using the exact model. Posterior mean phenological activity functions throughout the study period are depicted by the heatmap. The posterior mean peak activity day per year and corresponding 95% credible intervals are shown by black dots and horizontal lines. The linearized trend in peak activity days over time (obtained through OLS) and corresponding 95% credible interval is shown by a black line and white band across the study period. The posterior slope of change in peak activity day over the 49-year study period is displayed on top of the heatmap.

### 3.3 Joint species distribution modelling of soil-dwelling spider communities

Using the first-order Taylor approximation of our temporal disaggregation model enabled the estimation of a comprehensive joint species distribution model using temporally aggregated pitfall data for 331 species, yielding a wide array of results. Figure 4a shows the inferred phenological patterns for all spider species. Most species display a first phenological activity peak around April. Many of these species display a secondary activity peak in October. A limited number of species is predominantly active in winter. The phenological patterns for these species are clearly wrapped at the turn of the year, as enforced through the cyclic B-splines. For each species, differential activity patterns through time and space can be visualised (Fig. 4a). Upon linearizing the obtained inter-annual trends through ordinary least squares (at each posterior draw to propagate uncertainty), we obtain a summarised overview of linear trend slopes (Fig. 4b). We find strong statistical support for 82 species showing linear declines, and for 73 species showing linear increases. We could find strong statistical support for any linear trend for the remaining species, which can, for instance be due to insufficient data or non-linear trend patterns. The model also provides estimates on the influence of provided covariates for all species (Fig. 4c). By relying on a latent factor approach to model residual association patterns among sites and species, our model constitutes a model-based ordination technique that accounts for phenological influences in temporally aggregated data (Fig. 4d).

**Figure 4.**
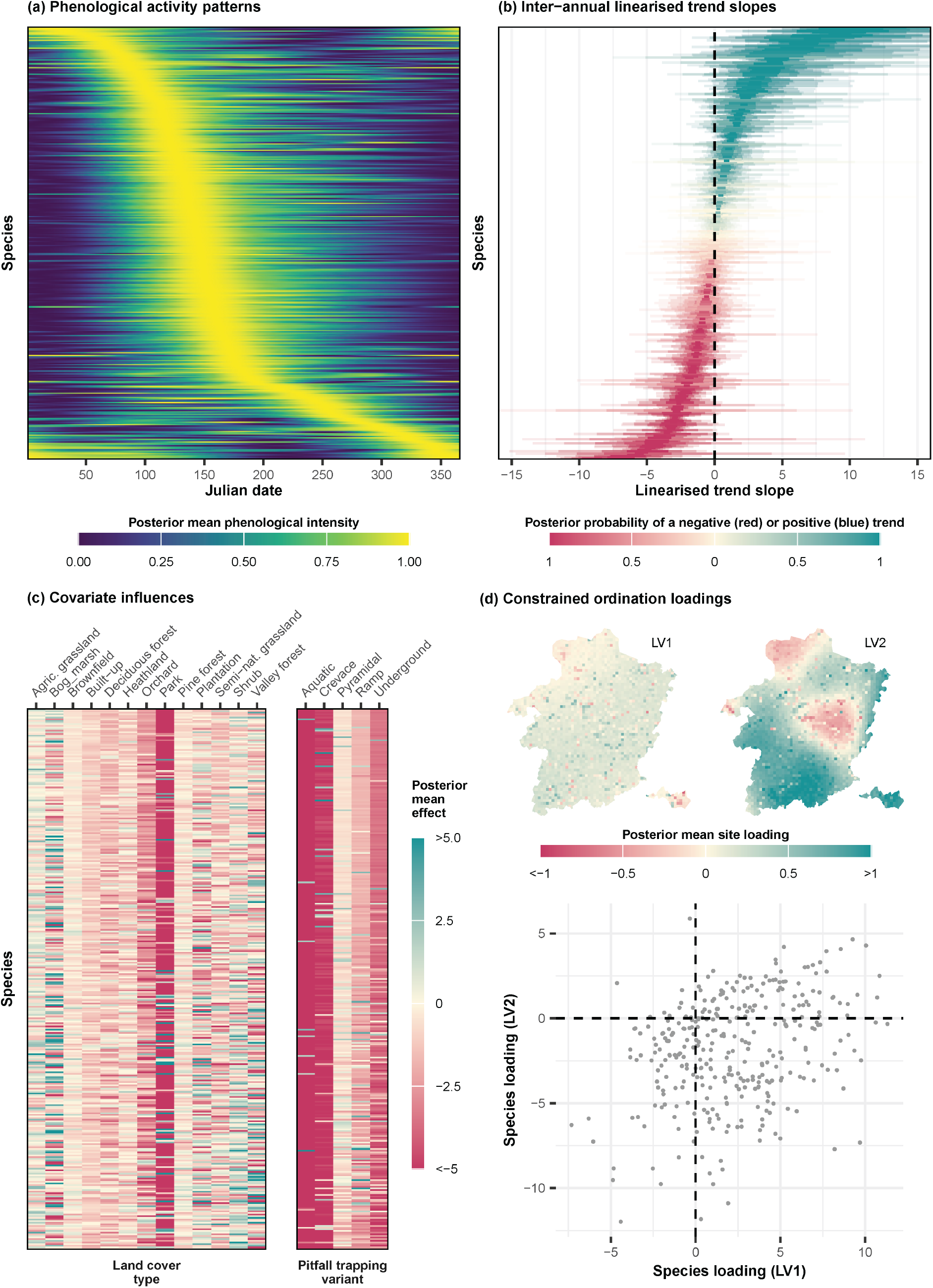
Condensed output of the joint species distribution model for pitfall trapping data. (a) Posterior mean phenological activity functions for all included species, with species ordered by the posterior mean peak activity day. For each species, the posterior mean B-spline projected Gaussian process capturing phenology is rescaled between 0 and 1. (b) Posterior linearised trend slopes for all included species, with species ordered by the posterior mean trend slopes. 50%, 80%, 95% and 99% credible intervals are depicted by varying transparency. (c) Posterior mean regression coefficients for included covariates and for all included species. Both land cover type related and pitfall trapping variant covariates are included. The dominant land cover, arable land, and the most frequently used variant, the conventional pitfall trap, constitute the reference levels in the analyses, to which all other effects should be compared with. (d) Posterior mean site and species loadings obtained from the latent factor ordination component in the model, depicting loadings along the two primary latent variables. Site loadings are geographically represented on a map of the study area, while species loadings are represented using a dot plot, with each dot representing a single species. Posterior mean loadings were computed using the first MCMC chain only, due to the possible sign symmetries among chains.

## 4 Discussion

The integrated and retrospective analysis of biodiversity data is essential to understand temporal patterns, to address shifting baselines and to inform present-day management practices (Magurran et al., 2010; Kindsvater et al., 2018; Fajgenblat et al., 2025). These efforts often require the combined analysis of data sources that do not necessarily match in the way they have been collected (Gotelli and Colwell, 2001; Magurran et al., 2010). Even though trapping techniques are relatively well standardised in themselves, they yield counts that are aggregated over sampling intervals that are often temporally misaligned, which can be problematic in the presence of phenological patterns. Although trapping data are abundantly available in ecology, the issue of aggregated count data in the face of phenological patterns has received remarkably little attention. In this paper, we outline a temporal disaggregation approach for trapping data in ecology. In addition to an exact model derived from the data generating process underlying temporal aggregation, we also provide computationally efficient Taylor approximations. We extend earlier work by Kotze et al. (2012) by estimating phenology more flexibly, by explicitly acknowledging the temporal aggregation process and how this affects phenological estimation, and by allowing arbitrarily overlapping sampling intervals, paving the way for the integrated analysis of retrospectively available trapping datasets.

Through simulations and through our case study, we show that failing to account for phenological influences and temporal disaggregation results in poor inference on quantities of interest, even if they are unrelated to phenology. While the exact model displays the best performance to deal with temporally aggregated data, it is computationally expensive, with costs increasing along sampling duration. When inferring phenological patterns is only of secondary importance, the results of our simulations and case study indicate that the use of the first-order Taylor series approximation is sufficient, and that the more complex approaches do not offer discernible advantages when inferring quantities unrelated to phenology or when optimizing predictive performance.

When inferring phenology or phenological change is a primary goal, our simulations indicate and our case study suggests that the exact model outperforms the approximate models. In our simulation study, the second-order Taylor approximation matches the performance of the exact model in several circumstances, and also provides significant advantages in inferring phenological change compared to the first-order Taylor approximation. The differences between the three methods were, however, relatively small in the case study involving real data.

Despite its simplicity, we find the first-order Taylor approximation to perform relatively well. Even though the second-order Taylor approximation and the exact model tend to outperform it when the primary interest lies in quantifying phenology, the gains are relatively modest. Conveniently, the first-order Taylor approximation can readily be fitted using conventional GAM fitting routines, such as the gam function of the mgcv package in R, by specifying the sampling duration of each sampling event through a logarithmic offset, and by defining a smoothing spline over the midpoint of the sampling interval. We therefore recommend this approach to most applied researchers. The use of the second-order Taylor approximation and the exact model requires bespoke modelling, e.g. through probabilistic programming languages such as Stan or Nimble. Especially when inferring phenology is of primary importance, when sampling intervals are long or when the phenological patterns tend to be rapidly changing, we recommend opting for these more complex approaches, for instance by using the scripts released along with this paper.

The benefits of using the exact or second-order Taylor approximation model to infer phenology can be expected to drop as the aleatoric uncertainty related to the true, underlying phenological activity function increases. If the underlying function is highly noisy, or if it varies across space and time, the aleatoric uncertainty can easily swamp the gains provided by the more exact models. In our case study, we accounted for yearly varying phenological activity functions, but we assumed phenology to be constant across space. While this might have been a sensible assumption given the relatively small study area, the outlined approach in Section 2.2.3 provides opportunities to account for such patterns through further extensions. Accounting for trapping fatigue or digging-in effects, i.e. a changing trapping efficiency throughout the sampling interval (e.g. due to depletion of source individuals, trap saturation, flooding by rain, scavenging, disturbance by rooting behaviour, temporary disturbance, …; Luff 1975; Digweed et al. 1995), constitutes an additional unexplored opportunity to further improve realism. The exact model in particular provides opportunities to account for such phenomena, and to estimate typical trapping fatigue patterns for each species. Even though meteorological conditions can influence trapping efficiency (Saska et al., 2013), we did not include them as predictors because our goal was to characterise the observed phenological activity pattern, regardless of its underlying drivers. Including meteorological variables would have removed part of the genuine phenological signal, namely the proximate, weather-induced variation that contributes to the realised activity pattern. Using the exact model, including daily meteorological predictors would constitute a straightforward extension. The first- and second-order Taylor approximation models, however, can only account for meteorological conditions summarised over the sampling interval, namely the mean in the first-order case, and the mean and variance in the second-order case.

Through our case study on soil-dwelling spiders in Limburg, we demonstrated how large historical datasets can yield rich output to answer ecological research questions on phenology, phenological change, inter-annual trends, and spatial patterns, by properly accounting for phenological patterns in temporally aggregated data. It also underscores the importance of properly recording the deployment start and end data, as this information is crucial for such applications. When analysing multi-species datasets, accounting for phenology in combination with model-based ordination approaches has recently been advocated to avoid spurious interspecific associations that merely arise from correlated phenological patterns (Lai, 2025). As pitfall trapping constitutes a frequently used method in community ecology, we here show how heterogeneities across datasets can be addressed to extend the spatial and temporal extent and resolution of such community analyses. The use of smoothing splines, Gaussian processes or other model-based approaches also overcomes limitations of simpler methods used to infer phenological change, such as the day of first sighting (Moussus et al., 2010). Our approach also paves the way for implementing more cost-effective biotic monitoring, as many taxa targeted by passive trapping techniques constitute excellent bioindicators (Rainio and Niemela, 2003). By statistically accounting for seasonal patterns rather than requiring year-round trapping, trapping effort can substantially be reduced and scheduled more flexibly.

The proposed interval-integrated B-spline approach we present is flexible and can be applied to any temporal resolution or extent. For instance, it could also be applied at an hourly resolution to resolve diurnal patterns in data that has been aggregated over multiple hours. Beyond the temporal dimension, this approach could be extended to the issue of spatial (dis)aggregation, which would not only find use in the field of ecology, but also in spatial epidemiology.

## Acknowledgements

We thank the Limburgse Koepel voor Natuurstudie (LIKONA) for providing the pitfall trapping data that inspired the development of the model and were used in the case studies. In particular, we are grateful to Marc Janssen for identifying the vast majority of collected spiders, Luc Crevecoeur for leading our collaborative efforts and preparing the data, and all other members of LIKONA’s Invertebrates Working Group for their valuable feedback during model development. T.N. gratefully acknowledges funding by Research Foundation Flanders (FWO, grant number G0A4121N). The computational resources and services used in this work were provided by the VSC (Flemish Supercomputer Center), funded by the Research Foundation Flanders (FWO) and the Flemish Government.

## References

Adis, J. (1979). Problems of interpreting arthropod sampling with pitfall traps. Zoologischer Anzeiger, 202(3/4):177–184.

Baars, M. A. (1979). Catches in pitfall traps in relation to mean densities of carabid beetles. Oecologia, 41(1):25–46.

Betancourt, M. (2018). A Conceptual Introduction to Hamiltonian Monte Carlo.

Bhattacharya, A. and Dunson, D. B. (2011). Sparse Bayesian infinite factor models. Biometrika, 98(2):291–306.

Bonte, D., Criel, P., Van Thournout, I., and Maelfait, J. P. (2003). Regional and local variation of spider assemblages (Araneae) from coastal grey dunes along the North Sea. Journal of Biogeography, 30(6):901–911.

Bowler, D. E., Boyd, R. J., Callaghan, C. T., Robinson, R. A., Isaac, N. J., and Pocock, M. J. (2025). Treating gaps and biases in biodiversity data as a missing data problem. Biological Reviews, 100(1):50–67.

Brown, G. R. and Matthews, I. M. (2016). A review of extensive variation in the design of pitfall traps and a proposal for a standard pitfall trap design for monitoring ground-active arthropod biodiversity. Ecology and Evolution, 6(12):3953–3964.

Bush, E. R., Abernethy, K. A., Jeffery, K., Tutin, C., White, L., Dimoto, E., Dikangadissi, J. T., Jump, A. S., and Bunnefeld, N. (2017). Fourier analysis to detect phenological cycles using long-term tropical field data and simulations. Methods in Ecology and Evolution, 8(5):530–540.

Carpenter, B., Gelman, A., Hoffman, M. D., Lee, D., Goodrich, B., Betancourt, M., Brubaker, M. A., Guo, J., Li, P., and Riddell, A. (2017). Stan: A probabilistic programming language. Journal of Statistical Software, 76(1).

Connolly, R., Schirmer, J., and Dunn, P. (1998). A daily rainfall disaggregation model. Agricultural and Forest Meteorology, 92(2):105–117.

De Saeger, S., Guelinckx, R., Oosterlynck, P., De Bruyn, A., Debusschere, K., Dhaluin, P., Erens, E., Hendrickx, P., Hennebel, D., Jacobs, I., Kumpen, M., Opdebeeck, J., Spanhove, T., Tamsyn, W., Van Oost, F., Van Dam, G., Van Hove, M., Wils, C., and Paelinckx, D. (2020). Biologische Waarderingskaart en Natura 2000 Habitatkaart, uitgave 2020. Technical report, Brussels.

Den Boer, P. J. (1979). The individual behaviour and population dynamics of some carabid beetles of forest. In den Boer, PJ., Thiele, HU., and Weber, F., editors, On the Evolution of Behaviour in Carabid Beetles, pages 151–166.

Desender, K. and Maelfait, J.-P. (1986). Pitfall trapping within enclosures: A method for estimating the relationship between the abundances of coexisting carabid species (Coleoptera: Carabidae). Ecography, 9(4):245–250.

Diggle, P. J., Moraga, P., Rowlingson, B., and Taylor, B. M. (2013). Spatial and spatiotemporal log-gaussian cox processes: Extending the geostatistical paradigm. Statistical Science, 28(4):542–563.

Digweed, SC., Currie, C. R., Cárcamo, H. A., and Spence, J. R. (1995). Digging out the “digging-in effect” of pitfall traps: Influences of depletion and disturbance on catches of ground beetles (Coleoptera: Carabidae). Pedobiologia, 39(6):561–576.

Eilers, P. H. C. and Marx, B. D. (1996). Flexible Smoothing with B-splines and Penalties. Statistical Science, 11(2):89–121.

Engel, J., Hertzog, L., Tiede, J., Wagg, C., Ebeling, A., Briesen, H., and Weisser, W. W. (2017). Pitfall trap sampling bias depends on body mass, temperature, and trap number: Insights from an individual-based model. Ecosphere, 8(4).

Fajgenblat, M., Wijns, R., De Knijf, G., Stoks, R., Lemmens, P., Herremans, M., Vanormelingen, P., Neyens, T., and De Meester, L. (2025). Leveraging Massive Opportunistically Collected Datasets to Study Species Communities in Space and Time. Ecology Letters, 28(3):e70094.

Ghysels, E., Santa-Clara, P., and Valkanov, R. (2004). The MIDAS Touch: Mixed Data Sampling Regression Models. Technical report.

Gotelli, N. J. and Colwell, R. K. (2001). Quantifying biodiversity: Procedures and pitfalls in the measurement and comparison of species richness. Ecology Letters, 4(4):379–391.

Greenslade, P. (1964). Pitfall Trapping as a Method for Studying Populations of Carabidae (Coleoptera). The Journal of Animal Ecology, 33:301–301.

Hallmann, C. A., Sorg, M., Jongejans, E., Siepel, H., Hofland, N., Schwan, H., Stenmans, W., Müller, A., Sumser, H., Hörren, T., Goulson, D., and De Kroon, H. (2017). More than 75 percent decline over 27 years in total flying insect biomass in protected areas. PLoS ONE, 12(10).

Hastie, T. and Tibshirani, R. (1986). Generalized Additive Models. Statistical Science, 3(1):297–318.

Hefley, T. J., Broms, K. M., Brost, B. M., Buderman, F. E., Kay, S. L., Scharf, H. R., Tipton, J. R., Williams, P. J., and Hooten, M. B. (2017). The basis function approach for modeling autocorrelation in ecological data. Ecology, 98(3):632–646.

Hernández-Carrasco, D., Tylianakis, J. M., Lytle, D. A., and Tonkin, J. D. (2025). Ecological and evolutionary consequences of changing seasonality. Science, 388(6750).

Heydemann, B. (1953). Agrarökologische Problematik.

Hilbe, J. M. (2011). Negative Binomial Regression. Cambridge University Press.

Hui, F. K., Taskinen, S., Pledger, S., Foster, S. D., and Warton, D. I. (2015). Model-based approaches to unconstrained ordination. Methods in Ecology and Evolution, 6(4):399– 411.

Johnson, O., Diggle, P., and Giorgi, E. (2019). A spatially discrete approximation to log-Gaussian Cox processes for modelling aggregated disease count data. Statistics in Medicine, 38(24):4871–4887.

Keil, P., Belmaker, J., Wilson, A. M., Unitt, P., and Jetz, W. (2013). Downscaling of species distribution models: A hierarchical approach. Methods in Ecology and Evolution, 4(1):82–94.

Kindsvater, H. K., Dulvy, N. K., Horswill, C., Juan-Jordá, M. J., Mangel, M., and Matthiopoulos, J. (2018). Overcoming the Data Crisis in Biodiversity Conservation. Trends in Ecology and Evolution, 33(9):676–688.

Kotze, D. J., O’Hara, R. B., and Lehvävirta, S. (2012). Dealing with varying detection probability, unequal sample sizes and clumped distributions in count data. PLoS ONE, 7(7).

Lai, H. R. (2025). Model-based ordination for phenological studies: From controlling sampling bias to inferring temporal associations. Methods in Ecology and Evolution, 16(7).

Lang, S. and Brezger, A. (2004). Bayesian P-Splines. Journal of Computational and Graphical Statistics, 13(1):183–212.

Lawton, J. H., Bignell, D. E., Bolton, B., Bloemers, G. F., Eggleton, P., Hammond, P. M., Hodda, M., Holt, R. D., Larsen, T. B., Mawdsley, N. A., Stork, N. E., Srivastava, D. S., and Watt, A. D. (1998). Biodiversity inventories, indicator taxa and effects of habitat modification in tropical forest. Nature, 391(6662):72–76.

Little, R. and Rubin, D. (2019). Statistical Analysis with Missing Data, Third Edition. Wiley.

Longcore, T. (2003). Terrestrial arthropods as indicators of ecological restoration success in coastal sage scrub (California, U.S.A.). Restoration Ecology, 11(4):397–409.

Luff, M. L. (1975). Some features influencing the efficiency of pitfall traps. Oecologia, 19(4):345–357.

Magurran, A. E., Baillie, S. R., Buckland, S. T., Dick, J. M. P., Elston, D. A., Scott, E. M., Smith, R. I., Somerfield, P. J., and Watt, A. D. (2010). Long-term datasets in biodiversity research and monitoring: Assessing change in ecological communities through time. Trends in Ecology and Evolution, 25(10):574–582.

Matechou, E., Dennis, E. B., Freeman, S. N., and Brereton, T. (2014). Monitoring abundance and phenology in (multivoltine) butterfly species: A novel mixture model. Journal of Applied Ecology, 51(3):766–775.

Miller, D. L. (2025). Bayesian views of generalized additive modelling. Methods in Ecology and Evolution, 16(3):446–455.

Miller-Rushing, A. J., Inouye, D. W., and Primack, R. B. (2008). How well do first flowering dates measure plant responses to climate change? The effects of population size and sampling frequency. Journal of Ecology, 96(6):1289–1296.

Monod, M., Blenkinsop, A., Brizzi, A., Chen, Y., Cardoso Correia Perello, C., Jogarah, V., Wang, Y., Flaxman, S., Bhatt, S., and Ratmann, O. (2023). Regularised B-splines Projected Gaussian Process Priors to Estimate Time-trends in Age-specific COVID-19 Deaths. Bayesian Analysis, 18(3):957–987.

Montgomery, G. A., Belitz, M. W., Guralnick, R. P., and Tingley, M. W. (2021). Standards and Best Practices for Monitoring and Benchmarking Insects. Frontiers in Ecology and Evolution, 8.

Moussus, J.-P., Julliard, R., and Jiguet, F. (2010). Featuring 10 phenological estimators using simulated data. Methods in Ecology and Evolution, 1(2):140–150.

Murphy, K. J., Ciuti, S., Burkitt, T., and Morera-Pujol, V. (2023). Bayesian areal disaggregation regression to predict wildlife distribution and relative density with low-resolution data. Ecological Applications, 33(8).

Norberg, A., Abrego, N., Blanchet, F. G., Adler, F. R., Anderson, B. J., Anttila, J., Araújo, M. B., Dallas, T., Dunson, D., Elith, J., Foster, S. D., Fox, R., Franklin, J., Godsoe, W., Guisan, A., O’Hara, B., Hill, N. A., Holt, R. D., Hui, F. K., Husby, M., Kålås, J. A., Lehikoinen, A., Luoto, M., Mod, H. K., Newell, G., Renner, I., Roslin, T., Soininen, J., Thuiller, W., Vanhatalo, J., Warton, D., White, M., Zimmermann, N. E., Gravel, D., and Ovaskainen, O. (2019). A comprehensive evaluation of predictive performance of 33 species distribution models at species and community levels. Ecological Monographs, 89(3).

O’Hara, R. B. and Kotze, D. J. (2010). Do not log-transform count data. Methods in Ecology and Evolution, 1(2):118–122.

Ovaskainen, O. and Soininen, J. (2011). Making more out of sparse data: Hierarchical modeling of species communities. Ecology, 92(2):289–295.

Ovaskainen, O., Tikhonov, G., Norberg, A., Guillaume Blanchet, F., Duan, L., Dunson, D., Roslin, T., and Abrego, N. (2017). How to make more out of community data? A conceptual framework and its implementation as models and software. Ecology Letters, 20(5):561–576.

Panaretos, V. M. and Zemel, Y. (2019). Statistical Aspects of Wasserstein Distances. Annual Review of Statistics and Its Application, 6(1):405–431.

Parmesan, C. (2007). Influences of species, latitudes and methodologies on estimates of phenological response to global warming. Global Change Biology, 13(9):1860–1872.

Pedersen, E. J., Miller, D. L., Simpson, G. L., and Ross, N. (2019). Hierarchical generalized additive models in ecology: An introduction with mgcv. PeerJ, 2019(5).

Peres-Neto, P. R., Olden, J. D., and Jackson, D. A. (2001). Environmentally constrained null models: Site suitability as occupancy criterion. Oikos, 93(1):110–120.

Popovic, G. C., Warton, D. I., Thomson, F. J., Hui, F. K., and Moles, A. T. (2019). Untangling direct species associations from indirect mediator species effects with graphical models. Methods in Ecology and Evolution, 10(9):1571–1583.

R Core Team (2023). R: A Language and Environment for Statistical Computing.

Rainio, J. and Niemela, J. (2003). Ground beetles (Coleoptera: Carabidae) as bioindicators. Biodiversity and Conservation, 12:487–506.

Rasmussen, C. E. and Williams, C. K. I. (2006). Gaussian Processes for Machine Learning. MIT Press.

Rubin, D. B. (1976). Inference and missing data. Biometrika, 63(3):581–92.

Rutten, S., Neyens, T., Duarte, E., and Faes, C. (2025). A Bayesian Geoadditive Model for Spatial Disaggregation.

Saska, P., van der Werf, W., Hemerik, L., Luff, M. L., Hatten, T. D., and Honek, A. (2013). Temperature effects on pitfall catches of epigeal arthropods: A model and method for bias correction. Journal of Applied Ecology, 50(1):181–189.

Seibold, S., Gossner, M. M., Simons, N. K., Blüthgen, N., Müller, J., Ambarlı, D., Ammer, C., Bauhus, J., Fischer, M., Habel, J. C., Linsenmair, K. E., Nauss, T., Penone, C., Prati, D., Schall, P., Schulze, E. D., Vogt, J., Wöllauer, S., and Weisser, W. W. (2019). Arthropod decline in grasslands and forests is associated with landscape-level drivers. Nature, 574(7780):671–674.

Strebel, N., Kéry, M., Schaub, M., and Schmid, H. (2014). Studying phenology by flexible modelling of seasonal detectability peaks. Methods in Ecology and Evolution, 5(5):483– 490.

ter Braak, C. J. F. (1986). Canonical Correspondence Analysis: A New Eigenvector Technique for Multivariate Direct Gradient Analysis. Ecology, 67(5):1167–1179.

Tikhonov, G., Opedal, Ø. H., Abrego, N., Lehikoinen, A., de Jonge, M. M., Oksanen, J., and Ovaskainen, O. (2020). Joint species distribution modelling with the r-package Hmsc. Methods in Ecology and Evolution, 11(3):442–447.

van der Veen, B., Hui, F. K., Hovstad, K. A., Solbu, E. B., and O’Hara, R. B. (2021). Model-based ordination for species with unequal niche widths. Methods in Ecology and Evolution, 12(7):1288–1300.

Van Strien, A. J., Plantenga, W. F., Soldaat, L. L., Van Swaay, C. A., and WallisDeVries, M. F. (2008). Bias in phenology assessments based on first appearance data of butterflies. Oecologia, 156(1):227–235.

Vehtari, A., Gelman, A., and Gabry, J. (2017). Practical Bayesian model evaluation using leave-one-out cross-validation and WAIC. Statistics and Computing, 27(5):1413–1432.

Vehtari, A., Gelman, A., Simpson, D., Carpenter, B., and Burkner, P. C. (2021). Rank-Normalization, Folding, and Localization: An Improved (Formula presented) for Assessing Convergence of MCMC (with Discussion)*†. Bayesian Analysis, 16(2):667–718.

Ver Hoef, J. M. and Boveng, P. L. (2007). Quasi-poisson vs. negative binomial regression: How should we model overdispersed count data? Ecology, 88(11):2766–2772.

Wakefield, J. and Salway, R. (2001). A statistical framework for ecological and aggregate studies. Journal of the Royal Statistical Society. Series A: Statistics in Society, 164(1):119–137.

Ward, D. F., New, T. R., and Yen, A. L. (2001). Effects of pitfall trap spacing on the abundance, richness and composition of invertebrate catches. Journal of Insect Conservation, 5:47–53.

Warton, D. I., Blanchet, F. G., O’Hara, R. B., Ovaskainen, O., Taskinen, S., Walker, S. C., and Hui, F. K. (2015). So Many Variables: Joint Modeling in Community Ecology. Trends in Ecology and Evolution, 30(12):766–779.

Weiss, D. J., Lucas, T. C., Nguyen, M., Nandi, A. K., Bisanzio, D., Battle, K. E., Cameron, E., Twohig, K. A., Pfeffer, D. A., Rozier, J. A., Gibson, H. S., Rao, P. C., Casey, D., Bertozzi-Villa, A., Collins, E. L., Dalrymple, U., Gray, N., Harris, J. R., Howes, R. E., Kang, S. Y., Keddie, S. H., May, D., Rumisha, S., Thorn, M. P., Barber, R., Fullman, N., Huynh, C. K., Kulikoff, X., Kutz, M. J., Lopez, A. D., Mokdad, A. H., Naghavi, M., Nguyen, G., Shackelford, K. A., Vos, T., Wang, H., Smith, D. L., Lim, S. S., Murray, C.J., Bhatt, S., Hay, S. I., and Gething, P. W. (2019). Mapping the global prevalence, incidence, and mortality of Plasmodium falciparum, 2000–17: A spatial and temporal modelling study. The Lancet, 394(10195):322–331.

Wood, S. N. (2017). Generalized Additive Models. Chapman and Hall/CRC.

Zabel, F. and Poschlod, B. (2023). The Teddy tool v1.1: Temporal disaggregation of daily climate model data for climate impact analysis. Geoscientific Model Development, 16(18):5383–5399.

Zhang, L., Carpenter, B., Gelman, A., and Vehtari, A. (2022). Pathfinder: Parallel quasi-Newton variational inference. Journal of Machine Learning Research, 23:1–49.

Zipkin, E. F., Dewan, A., and Andrew Royle, J. (2009). Impacts of forest fragmentation on species richness: A hierarchical approach to community modelling. Journal of Applied Ecology, 46(4):815–822.

