## Supplementary Information for "Temporal disaggregation through interval-integrated B-splines for the integrated analysis of trapping counts in ecology"

### Supporting text

### S1 Extended methods

#### S1.1 Derivation of interval-integrated B-splines for Gaussian responses

Let $Y_i$ denote the random variable representing the aggregate outcome during sampling event $i = 1, 2, \dots, n$. $Y_i$ can be decomposed into the sum of (unobserved) daily outcomes: $Y_i = Z_{i,d_{\text{start}(i)}} + \dots + Z_{i,d_{\text{end}(i)}}$, where $Z_{i,d}$ denotes the random variable for the number of individuals during day $d$ for the $i$-th sampling event. We assume that $Z_{i,d}$ follows a Gaussian distribution:

$$Z_{i,d} \sim \text{Normal}\left(\mu_i + f(d), \sigma^2\right), \tag{1}$$

where $\mu_{i,d}$ is a parametric sample-specific linear predictor part, $f(d)$ is a non-parametric day-specific phenological part, and $\sigma$ is the daily noise parameter. We assume the phenological activity function $f(d)$ can be described using cubic B-splines, as the weighted sum of an appropriate number of $k$ basis functions:

$$f(d) = \sum_{j=1}^{k} B_j(d)\gamma_j = \boldsymbol{B}_d^\intercal \boldsymbol{\gamma}, \tag{2}$$

where $B_j(d)$ is the $j$-th basis function and $\gamma_j$ is the corresponding basis function weight.

We assume that the daily outcomes $Z_{i,d}$ are conditionally independent. The sum $Y_i = Z_{i,d_{\text{start}(i)}} + \dots + Z_{i,d_{\text{end}(i)}}$ of independent normally distributed random variables can be written as:

$$Y_i \sim \text{Normal}\left(\sum_{d=d_{\text{start}(i)}}^{d_{\text{end}(i)}} (\mu_i + f(d)), \sum_{d=d_{\text{start}(i)}}^{d_{\text{end}(i)}} \sigma^2\right). \tag{3}$$

Defining $m_i = d_{\text{end}(i)} - d_{\text{start}(i)} + 1$ as the number of days comprised in the sampling interval of sampling event $i$, we have:

$$\begin{aligned}\sum_{d=d_{\text{start}(i)}}^{d_{\text{end}(i)}} (\mu_i + f(d)) &= m_i\mu_i + \sum_{d=d_{\text{start}(i)}}^{d_{\text{end}(i)}} \sum_{j=1}^{k} B_j(d)\gamma_j \\ &= m_i\mu_i + \sum_{j=1}^{k}\left(\gamma_j \sum_{d=d_{\text{start}(i)}}^{d_{\text{end}}(i)} B_j(d)\right).\end{aligned} \tag{4}$$

Conveniently, $\sum_{d=d_{\text{start}}}^{d_{\text{end}}} B_j(d)$ represents the sum of the $j$-th basis function evaluated over all days $\{d_{\text{start}}, \ldots, d_{\text{end}}\}$, which can be pre-computed for each sampling event $i$ prior to model estimation. Since these modified basis functions are Riemann sums over the full temporal interval covered by sampling event $i$, we coin them interval-integrated basis functions, and notationally abbreviate them as follows:

$$B^*_{i,j} = \sum_{d=d_{\text{start}(i)}}^{d_{\text{end}}(i)} B_j(d). \tag{5}$$

Hence, upon pre-computing the integral-integrated basis functions $\boldsymbol{B}^*_{\boldsymbol{i}}$ (Listing 1), the model simplifies to:

$$Y_i \sim \text{Normal}\left(m_i\mu_i + \boldsymbol{B}_{\boldsymbol{i}}^{*\intercal}\boldsymbol{\gamma}, m_i\sigma^2\right). \tag{6}$$

```
# Load required libraries
library(mgcv)

# Function to generate day sequences using day of the year indices
generate_day_sequence <- function(start_day, end_day) {
  if (start_day <= end_day) {
    # Simple case: start day is before end day within the same year
    return(seq(start_day, end_day))
  } else {
    # Wraparound case: part of the sequence is before and part after
        year-end
    return(c(seq(1, end_day), seq(start_day, 366)))
```

```
  }
}

# Function to compute interval-integrated basis functions
iibf <- function(start_days, end_days, N_basis_functions = 12, ord = 4) {
  # Compute daily basis functions with cyclic spline
  daily_basis_functions <- cSplineDes(1:366, seq(1, 366, length.out =
      N_basis_functions), ord = 4)
  # Generate list of day sequences for each sampling event
  day_sequences <- mapply(generate_day_sequence, start_days, end_days,
      SIMPLIFY = F)
  # Compute interval-integrated basis functions
  iibf_list <- lapply(day_sequences, function(day_sequence)
      colSums(daily_basis_functions[day_sequence,]))
  return(do.call(rbind, iibf_list))
}

# Example computation for 100 randomly sampled start and end days
iibf_design_matrix <- iibf(start_days = sample(1:366, 100), end_days =
    sample(1:366, 100), N_basis_functions = 12)
```

Listing 1: R code for the computation of interval-integrated basis functions, which can be supplied as a design matrix for subsequent modelling.

#### S1.2 Derivation of interval-integrated B-splines for count responses

As developed in the main text, the exact linear predictor for aggregate count data corresponds to:

$$
\begin{aligned}
&\log\left(\lambda_{i,d_{\text{start}(i)}} + \ldots + \lambda_{i,d_{\text{end}(i)}}\right) \\
&= \log\left(\exp\left(\mu_i + f(d_{\text{start}(i)})\right) + \ldots + \exp\left(\mu_i + f(d_{\text{end}(i)})\right)\right) \\
&= \mu_i + \log\left(\exp\left(f(d_{\text{start}(i)})\right) + \ldots + \exp\left(f(d_{\text{end}(i)})\right)\right) \\
&= \mu_i + \log\left(\exp\left(\boldsymbol{B}_{d_{\text{start}}}^{\intercal}\gamma\right) + \ldots + \exp\left(\boldsymbol{B}_{d_{\text{end}}}^{\intercal}\gamma\right)\right). \qquad (7)
\end{aligned}
$$

In contrast to the Gaussian case, this expression cannot be further simplified by substitution of pre-computed interval-integrated basis functions, and the computation of the sum of daily exponentials for each sampling event can be prohibitive for moderate to large datasets. A computationally efficient interval-integrated approximation of the exponential B-splines can, however, be achieved through a first- or second-order Taylor series expansion around the midpoint $d^*$ of the sampling interval.

Let $g(d) = \exp(f(d))$ be the exponentiated phenological B-spline function. The function $g(d)$ can be approximated using a second-order Taylor approximation as follows:

$$g(d) \approx g(d^*) + \frac{g'(d^*)}{1!}(d - d^*) + \frac{g''(d^*)}{2!}(d - d^*)^2. \tag{8}$$

Through the chain rule, the exponentiated phenological value on day $d$ can therefore be reformulated in terms of the midpoint $d^*$ of the sampling interval:

$$\begin{aligned}\exp(f(d)) \approx & \exp(f(d^*)) + \exp(f(d^*))f'(d^*)(d - d^*) \\ & + \frac{\exp(f(d^*))\left((f'(d^*))^2 + f''(d^*)\right)(d - d^*)^2}{2}.\end{aligned} \tag{9}$$

This approximation can be used to substitute the sum of daily exponentials:

$$\begin{aligned}&\exp\left(f(d_{\text{start}(i)})\right) + \ldots + \exp\left(f(d_{\text{end}(i)})\right) \\ &\approx \sum_{d=d_{\text{start}(i)}}^{d_{\text{end}(i)}} \exp(f(d_i^*)) + \sum_{d=d_{\text{start}(i)}}^{d_{\text{end}(i)}} \exp(f(d_i^*))f'(d_i^*)(d - d_i^*) \\ &\quad + \sum_{d=d_{\text{start}(i)}}^{d_{\text{end}(i)}} \frac{\exp(f(d_i^*))\left((f'(d_i^*))^2 + f''(d_i^*)\right)(d - d_i^*)^2}{2}.\end{aligned} \tag{10}$$

Since $\sum_{d=d_{\text{start}(i)}}^{d_{\text{end}(i)}} \exp(f(d_i^*)) = m_i \exp(f(d_i^*))$ (with $m_i$ the duration of the sampling interval) and since $\sum_{d=d_{\text{start}(i)}}^{d_{\text{end}(i)}} \exp(f(d_i^*))f'(d_i^*)(d - d_i^*) = 0$ (as $d^*$ is the midpoint of the sampling interval), the sum of daily exponentials can further be simplified to:

$$\begin{aligned}&\exp\left(f(d_{\text{start}(i)})\right) + \ldots + \exp\left(f(d_{\text{end}(i)})\right) \\ &\approx m_i \exp(f(d_i^*)) + \exp(f(d_i^*))\left((f'(d_i^*))^2 + f''(d_i^*)\right)\frac{\sum_{d=d_{\text{start}(i)}}^{d_{\text{end}(i)}}(d - d_i^*)^2}{2} \\ &\approx m_i \exp(f(d_i^*))\left(1 + c_i\left((f'(d_i^*))^2 + f''(d_i^*)\right)\right) \\ &\approx m_i \exp(\boldsymbol{B}_{d_i^*}^{\mathsf{T}}\boldsymbol{\gamma})\left(1 + c_i\left(\left(\boldsymbol{B'}_{d_i^*}^{\mathsf{T}}\boldsymbol{\gamma}\right)^2 + \boldsymbol{B''}_{d_i^*}^{\mathsf{T}}\boldsymbol{\gamma}\right)\right),\end{aligned} \tag{11}$$

where $c_i = \frac{\sum_{d=d_{\text{start}(i)}}^{d_{\text{end}(i)}}(d-d_i^*)^2}{2m_i}$ is a constant that can be precomputed for each sampling event $i$. The computation of $f(d_i^*)$ and its derivatives $f'(d_i^*)$ and $f''(d_i^*)$ is straightforward due to the properties of B-splines. These correspond to $\boldsymbol{B}_{d_i^*}^{\mathsf{T}}\boldsymbol{\gamma}$, $\boldsymbol{B'}_{d_i^*}^{\mathsf{T}}\boldsymbol{\gamma}$ and $\boldsymbol{B''}_{d_i^*}^{\mathsf{T}}\boldsymbol{\gamma}$, respectively, where $\boldsymbol{B}$, $\boldsymbol{B}'$ and $\boldsymbol{B}''$ can be provided as design matrices. Hence:

$$\begin{aligned}&\exp\left(f(d_{\text{start}(i)})\right) + \ldots + \exp\left(f(d_{\text{end}(i)})\right)\\&\approx m_i \exp(\boldsymbol{B}^{\intercal}_{d_i^*}\boldsymbol{\gamma}) \left(1 + c_i \left(\left(\boldsymbol{B}'^{\intercal}_{d_i^*}\boldsymbol{\gamma}\right)^2 + \boldsymbol{B}''^{\intercal}_{d_i^*}\boldsymbol{\gamma}\right)\right).\end{aligned} \tag{12}$$

The linear predictor in Eq. 7 can therefore be approximated as:

$$\begin{aligned}&\log\left(\lambda_{i,d_{\text{start}(i)}} + \ldots + \lambda_{i,d_{\text{end}(i)}}\right)\\&\approx \mu_i + \log(m_i) + \boldsymbol{B}^{\intercal}_{d_i^*}\boldsymbol{\gamma} + \log\left(1 + c_i \left(\left(\boldsymbol{B}'^{\intercal}_{d_i^*}\boldsymbol{\gamma}\right)^2 + \boldsymbol{B}''^{\intercal}_{d^*}\boldsymbol{\gamma}\right)\right).\end{aligned} \tag{13}$$

The structure of this linear predictor precludes the use of generic model fitting software, which is also the case for the exact model (Eq. 7). Implementing these models is, however, straightforward using probabilistic programming languages such as Stan.

A first-order Taylor approximation follows from an analogous derivation and leads to the following linear predictor:

$$\log\left(\lambda_{i,d_{\text{start}(i)}} + \ldots + \lambda_{i,d_{\text{end}(i)}}\right) \approx \mu_i + \log(m_i) + \boldsymbol{B}^{\intercal}_{d_i^*}\boldsymbol{\gamma}, \tag{14}$$

which can readily be implemented using conventional GAM fitting routines, such as the `gam` function of the `mgcv` package in R. This approach involves specifying the trapping interval duration as an offset and evaluating the phenological spline at the midpoint of the trapping interval (Listing 2).

```r
library(mgcv)
fit <- gam(aggregated_count ~ predictor_1 + ... + predictor_n +
             offset(log(duration)) + s(midpoint, bs = "cc"),
             family = "poisson", data = data)
```

Listing 2: R code for fitting the first-order Taylor approximation temporal disaggregation model for count data using mgcv, assuming a Poisson likelihood.

#### S1.3 Derivation of the compound negative binomial distribution

Here, we show that if the daily counts $Z_{i,d}$ are modelled as a Poisson–Gamma mixture, better known as the negative binomial distribution, then the aggregated count $Y_i = \sum_{d=d_{\text{start}(i)}}^{d_{\text{end}(i)}} Z_{i,d}$ is also negative binomially distributed with the same dispersion parameter.

Assume that each daily count is conditionally independent given a latent variable $u$, such that:

$$Z_{i,d} \mid u \sim \text{Poisson}(u \cdot \lambda_{i,d}) \quad \text{with} \quad u \sim \text{Gamma}(\phi, \phi), \tag{15}$$

so that $\text{E}(u) = 1$ and $\text{Var}(u) = 1/\phi$. Marginally, each $Z_{i,d}$ then follows a negative binomial distribution:

$$Z_{i,d} \sim \text{NegBin}(\lambda_{i,d}, \phi), \tag{16}$$

with mean $\text{E}(Z_{i,d}) = \lambda_{i,d}$ and variance $\text{Var}(Z_{i,d}) = \lambda_{i,d} + \frac{\lambda_{i,d}^2}{\phi}$. Given the conditional independence, the sum over the sampling interval satisfies:

$$Y_i \mid u \sim \text{Poisson}\left(u \cdot \sum_{d=d_{\text{start}(i)}}^{d_{\text{end}(i)}} \lambda_{i,d}\right). \tag{17}$$

Marginalizing over $u$ yields:

$$Y_i \sim \text{NegBin}\left(\sum_{d=d_{\text{start}(i)}}^{d_{\text{end}(i)}} \lambda_{i,d}, \phi\right), \tag{18}$$

demonstrating that the aggregated count retains the negative binomial form with a mean equal to the sum of the daily means and the dispersion parameter $\phi$ unchanged. This derivation justifies the extension of our temporal disaggregation model from a Poisson likelihood to a negative binomial likelihood.

#### S1.4 Extending the linear predictor for trapping data: methodological details

In the following, we provide additional methodological details for the univariate trapping model outlined in the main text.

The temporal random effects $\zeta_t$ are modelled smoothly by means of Gaussian processes, a powerful way of modelling time-series:

$$\boldsymbol{\zeta} \sim \mathcal{GP}\left(0, C^{\text{time}}\left(\Delta t\right)\right), \tag{19}$$

where $C^{\text{time}}\left(\cdot\right)$ is a covariance function that dictates how the covariance between years decays as a function of the time that separates them (Rasmussen and Williams, 2006). We specifically assume a species-specific exponentiated quadratic kernel as covariance function:

$$C^{\text{time}}\left(\Delta t\right) = \alpha^{\text{time}} \exp\left(-\frac{1}{2}\left(\frac{\Delta t}{\rho^{\text{time}}}\right)^2\right), \tag{20}$$

where $\alpha^{\text{time}}$ is the GP's marginal scale parameter and $\rho^{\text{time}}$ is the length scale parameter. The spatial random effects $\nu_s$ for each site $s$ are modelled as a linear combination of spatially correlated (i.e. structured) and uncorrelated (i.e. unstructured) random effects, weighted by the spatial signal $p$:

$$\nu_s = \left(1 - p\right)\nu_s^{\text{unstr}} + p\nu_s^{\text{str}}, \tag{21}$$

where $\nu_s^{\text{unstr}}$ and $\nu_s^{\text{str}}$ represent the spatially unstructured and structured random effects respectively. The spatially unstructured random effects are assumed to be normally distributed:

$$\nu_s^{\text{unstr}} \sim \text{Normal}\left(0, \alpha^{\text{space}}\right), \tag{22}$$

with $\alpha^{\text{space}}$ a scale parameter. Similarly to the temporal random effects, we use Gaussian processes to model the spatially structured random effects $\nu_s^{\text{str}}$. Exact Gaussian processes become impractical when being evaluated over more than a couple of hundred input locations due to their cubically scaling computational complexity. We use B-splines projected Gaussian processes (Monod et al., 2023) instead to facilitate the evaluation over a larger number of locations. B-splines projected Gaussian processes are similar to penalised splines but have been shown to outperform them (Monod et al., 2023). First, we define a two-dimensional tensor-product spline surface with $p$ basis functions along each dimension over the study area's square bounding box:

$$\nu_s^{\text{str}} = \sum_{k=1}^{p}\sum_{l=1}^{p}\left(w_{k,l} \cdot b_k\left(\text{lon}(s)\right) \cdot b_l\left(\text{lat}(s)\right)\right), \tag{23}$$

where $\text{lon}(s)$ and $\text{lat}(s)$ are the longitude and latitude of site$s$, $b_k\left(\cdot\right)$ and $b_l\left(\cdot\right)$ are the $k$'th and $l$'th cubic basis functions anchored to equally spaced knots along each dimension, and $w_{k,l}$ is their corresponding weight coefficient. Since most study areas do not match an exact square, some basis function products might equal zero (or some negligible value) across the entire study area. Accordingly, these basis functions combinations do not need to be evaluated and the corresponding weight coefficients $w_{k,l}$ do not need to be estimated, easing computation. The full set of B-spline weight coefficients $\boldsymbol{w}$ are modelled through an exact Gaussian process:

$$\boldsymbol{w} \sim \mathcal{GP}\left(0, C^{\text{space}}\left(\sqrt{(\Delta k)^2 + (\Delta l)^2}\right)\right), \tag{24}$$

where $C^{\text{space}}(\cdot)$ is a covariance function that dictates how the covariance between two weight coefficients decays as a function of the Euclidean distance between the knots to which their basis functions are anchored. In a similar way as for the temporal Gaussian process, we assume a latent dimension-specific exponentiated quadratic kernel as covariance function:

$$C^{\text{space}}\left(\sqrt{(\Delta k)^2 + (\Delta l)^2}\right) = \alpha^{\text{space}} \exp\left(-\frac{(\Delta k)^2 + (\Delta l)^2}{2(\rho^{\text{space}})^2}\right), \tag{25}$$

where $\rho^{\text{space}}$ is the length scale parameter. Note that our model only includes time and space in a separable way, and that it does not explicitly account for non-separable spatiotemporal effects. Whenever deemed relevant, spatiotemporal random effects can be included, e.g. through the inclusion of an anisotropic, three-dimensional Gaussian process.

#### S1.5 Prior specifications

##### S1.5.1 Single-species models

The following prior specifications pertain to all single-species models presented throughout the manuscript (`TDM_simple.stan`, `TDM_flexible.stan`, `TDM_yearlyphen.stan`). Note, however, that not all models include all mentioned parameters outlined below.

We assume weakly informative Normal$(0, 3)$ priors on the model intercept $\beta_0$ and regression coefficients $\boldsymbol{\beta}$. We use a moderately informative InvGamma$(5, 5)$ prior on the length scales of all Gaussian processes included in the models (i.e. $\rho^{\text{time}}$ for the interannual trend, $\rho^{\text{space}}$ for the spatial basis function weights and $\rho$ for the phenological basis function weights). Along with this specific prior, we scale the Gaussian processes' input values to the range $[-1, 1]$, to constrain the length scales to sensible values. We use a weakly informative Normal$^{+}(0, 3)$ prior on the Gaussian processes' marginal standard deviation. We use a uniform Uniform$(0, 1)$ prior on the spatial signal $p$. We use a weakly informative Normal$^+(0,3)$ on the scale of the spatial random effects $\alpha^{\text{space}}$. Specifically for the `TDM_yearlyphen.stan` model, the differences between historical and contemporary weights $\boldsymbol{\gamma}^{\text{diff}}$ are assumed to follow a normal distribution, with a weakly informative Normal$^+(0,1)$ prior on its scale parameter. The yearly deviations on the weights $\boldsymbol{\gamma}^{\text{iid}}$ are also assumed to follow a normal distribution, with a weakly informative Normal$^+(0,1)$ prior on its scale parameter. We use a weakly informative Normal$^+(0,3)$ prior for the negative binomial distribution's dispersion parameter $\phi$.

##### S1.5.2 Multi-species model

The priors for the multi-species model (`TDM_multispecies.stan`) are largely similar to the ones of the single-species models, with the following exceptions. The species-specific intercepts $\beta_{0,l}$ and regression coefficients $\boldsymbol{\beta_l}$ are modelled as normally distributed random effects, with a weakly informative Normal$^+(0,3)$ prior on the random effects' scale. We use a moderately informative InvGamma$(5,5)$ prior on the length scale and a weakly informative Normal$^+(0,3)$ prior on the marginal standard deviation of the Gaussian processes that are used to model the common phenological pattern across species $\boldsymbol{\gamma}^{\text{common}}$ and species-specific deviations $\boldsymbol{\gamma}^{\text{specific}}$. Following Bhattacharya and Dunson (2011), we use multiplicative gamma process shrinkage priors on the latent species loadings, with a Gamma$(2,1)$ prior on the first latent dimension and a Gamma$(6,1)$ prior on subsequent latent dimensions.

#### References


Bhattacharya, A. and Dunson, D. B. (2011). Sparse Bayesian infinite factor models. *Biometrika*, 98(2):291–306.

Monod, M., Blenkinsop, A., Brizzi, A., Chen, Y., Cardoso Correia Perello, C., Jogarah, V., Wang, Y., Flaxman, S., Bhatt, S., and Ratmann, O. (2023). Regularised B-splines Projected Gaussian Process Priors to Estimate Time-trends in Age-specific COVID-19 Deaths. *Bayesian Analysis*, 18(3):957–987.

Rasmussen, C. E. and Williams, C. K. I. (2006). *Gaussian Processes for Machine Learning.* MIT Press.

### Supporting figures

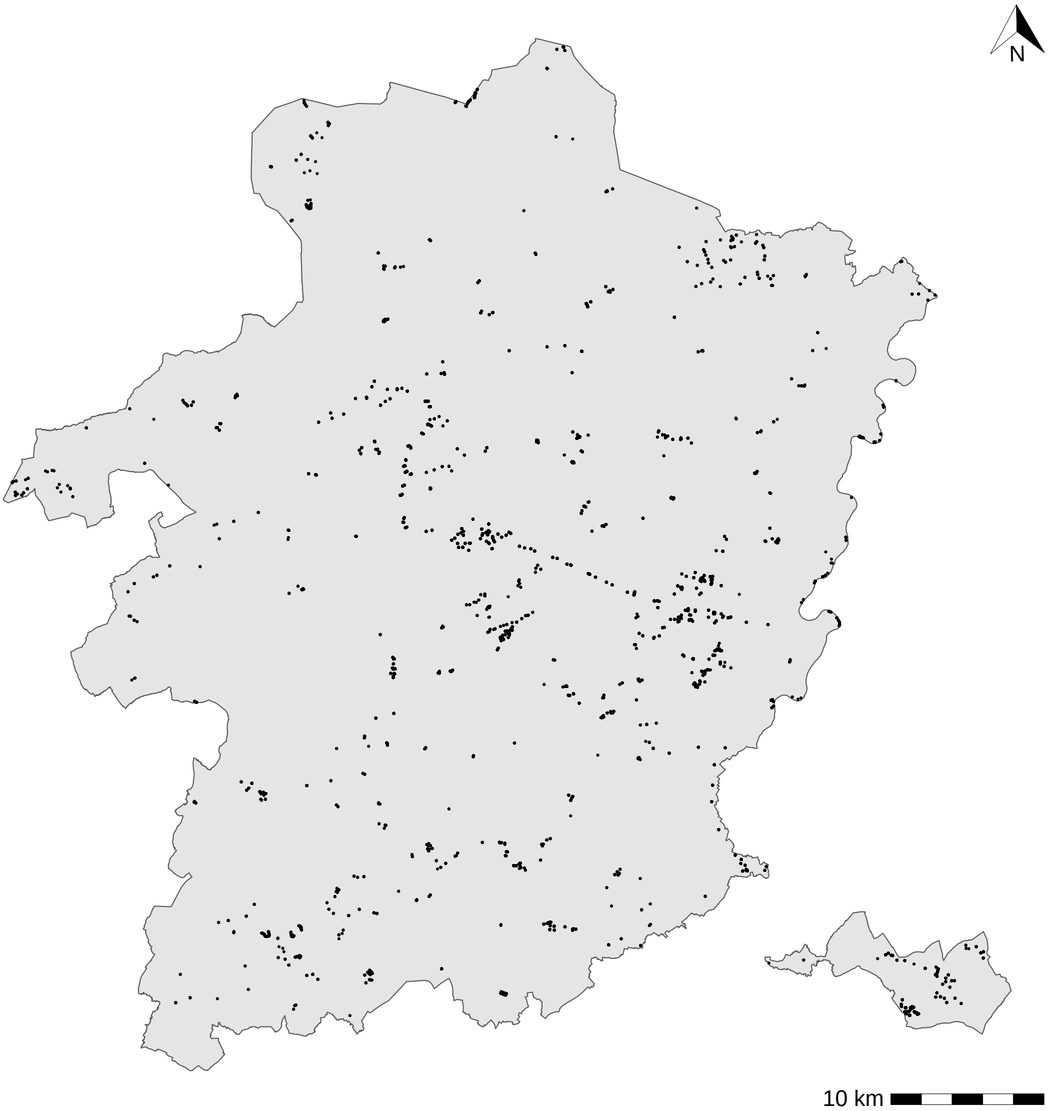


Figure 1: Overview of locations where pitfall traps were placed across the province of Limburg.

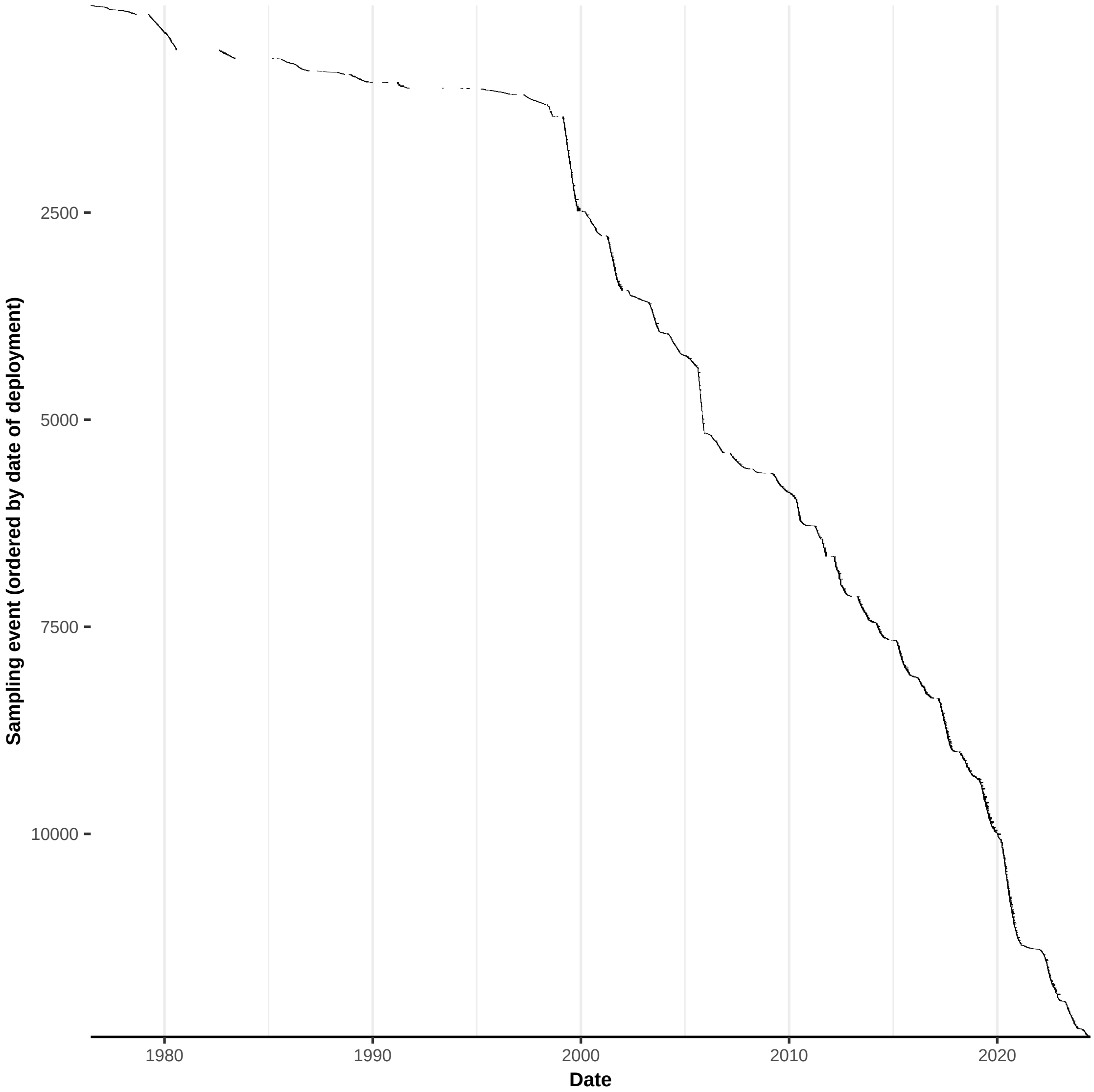


Figure 2: Overview of sampling intervals during which pitfall traps were placed across the study period.

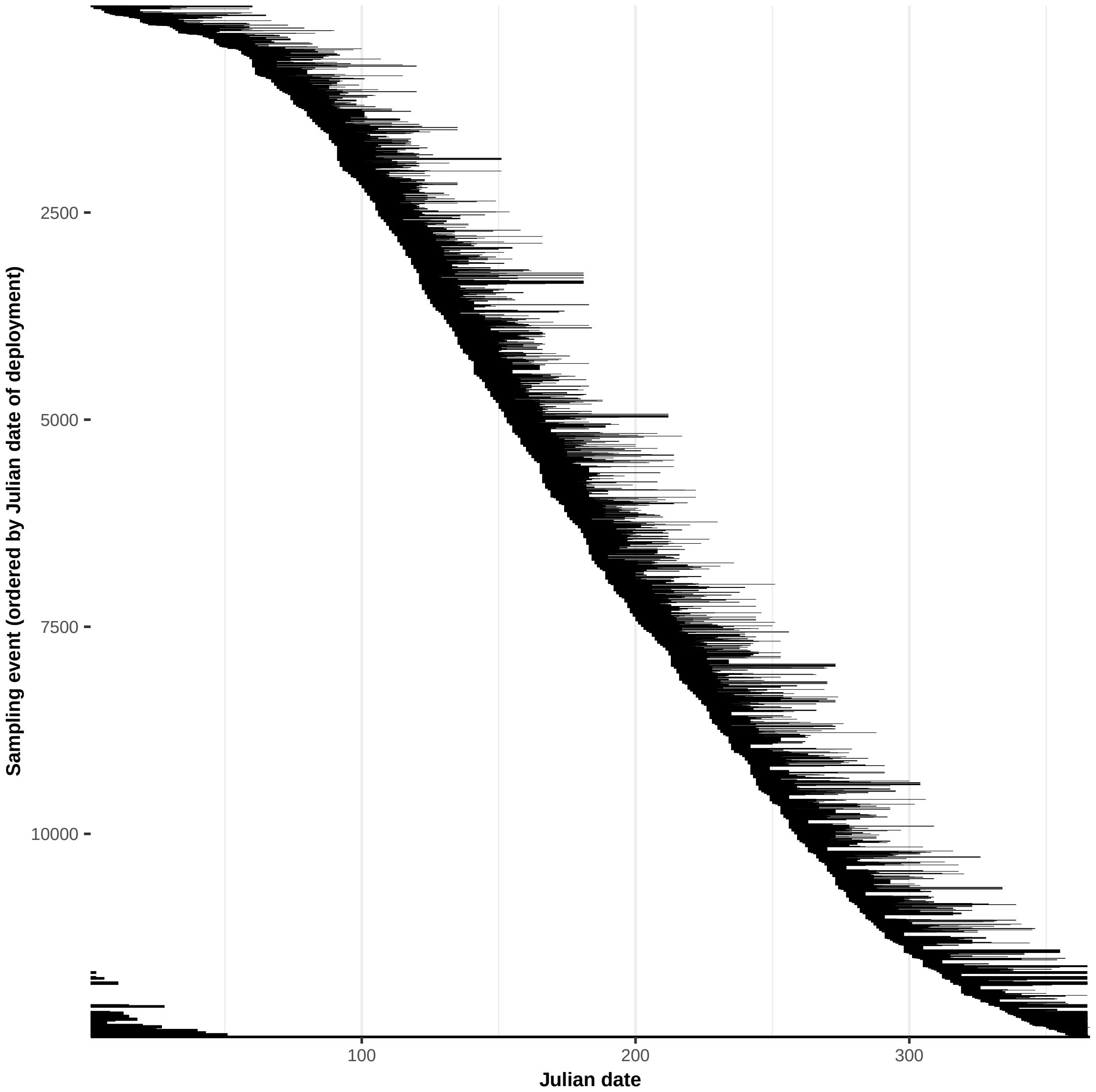


Figure 3: Overview of sampling intervals during which pitfall traps were placed across days of the year.
